# Structure of the human heparan-α-glucosaminide *N*-acetyltransferase (HGSNAT)

**DOI:** 10.1101/2023.10.23.563672

**Authors:** Vikas Navratna, Arvind Kumar, Jaimin K. Rana, Shyamal Mosalaganti

## Abstract

Degradation of heparan sulfate (HS), a glycosaminoglycan (GAG) comprised of repeating units of *N*-acetylglucosamine and glucuronic acid, begins in the cytosol and is completed in the lysosomes. Acetylation of the terminal non-reducing amino group of α-D-glucosamine of HS is essential for its complete breakdown into monosaccharides and free sulfate. Heparan-α-glucosaminide *N*-acetyltransferase (HGSNAT), a resident of the lysosomal membrane, catalyzes this essential acetylation reaction by accepting and transferring the acetyl group from cytosolic acetyl-CoA to terminal α-D-glucosamine of HS in the lysosomal lumen. Mutation-induced dysfunction in HGSNAT causes abnormal accumulation of HS within the lysosomes and leads to an autosomal recessive neurodegenerative lysosomal storage disorder called mucopolysaccharidosis IIIC (MPS IIIC). There are no approved drugs or treatment strategies to cure or manage the symptoms of, MPS IIIC. Here, we use cryo-electron microscopy (cryo-EM) to determine a high-resolution structure of the HGSNAT-acetyl-CoA complex, the first step in HGSNAT catalyzed acetyltransferase reaction. In addition, we map the known MPS IIIC mutations onto the structure and elucidate the molecular basis for mutation-induced HGSNAT dysfunction.

## Introduction

Degradation of the extracellular matrix proteoglycans begins in the cytosol, where they are proteolyzed to glycosaminoglycans (GAGs) such as heparan sulfate (HS), chondroitin sulfate, dermatan sulfate, and keratan sulfate. The proteolyzed GAGs are further degraded to monosaccharides and free sulfate residues within the lysosomes in GAG specific multi-enzyme degradation pathways. Dysfunction, often inherited, of the enzymes involved in GAG degradation causes abnormal accumulation of the partial degradation products within the lysosomes, resulting in mucopolysaccharidoses (MPSs), a collective term for the group of rare autosomal recessive lysosomal storage disorders characterized by the accumulation of partially degraded GAGs (Coutinho *et al*, 2012; Huizing & Gahl, 2020; Klein *et al*, 1978; Platt *et al*, 2018; Ruivo *et al*, 2009). HS is a GAG made of repeating units of *N*-acetylglucosamine and glucuronic acid, and impairment of enzymes in the HS degradation pathway causes mucopolysaccharidosis III (MPS III), or Sanfilippo’s syndrome that is characterized by abnormal storage of HS degradation pathway intermediates within the lysosomes in all organs and excretion of these intermediates in the urine. The onset of the symptoms of MPS III, which include organomegaly, abnormal joint mobility, compromised cardiac function, degeneration of vision and hearing, and dementia, often begins in juveniles and ends in premature death. Currently, no therapies for MPS III are available (Coutinho *et al,* 2012; Huizing & Gahl, 2020; McBride & Flanigan, 2021; Pshezhetsky *et al*, 2018).

Depending on the dysfunctional enzyme, MPS III is classified into four subtypes – MPS IIIA-IIID. Of these four MPS III subtypes, MPS IIIC (prevalence rate of 1 in 1.4 million) is caused by the dysfunction of heparan-α-glucosaminide *N*-acetyltransferase (HGSNAT, EC 2.3.1.78). HGSNAT is the only enzyme of the GAG pathway that is not a hydrolase. It catalyzes the only known biosynthetic reaction of the GAG degradation pathway within the lysosome, that is, the acetyl-CoA (ACO) mediated *N-*acetylation of the terminal non-reducing amino group of α-D-glucosamine (Fan *et al*, 2006; Hrebicek *et al*, 2006; Huizing & Gahl, 2020; Klein *et al,* 1978; Nagel *et al*, 2019; Pshezhetsky *et al,* 2018). Acetylation of the terminal α-D-glucosamine group is essential for subsequent HS degradation within the lysosomes. HGSNAT is not homologous to any known proteins, including acetyl-CoA binding proteins, *N*-acetyltransferases, and other lysosomal proteins. HGSNAT mRNA has two translation start sites (M1 and M29), yielding two functionally active isoforms (73 kDa and 70 kDa) of HGSNAT, both targeted to the lysosomal membrane (Durand *et al*, 2010; Fan *et al*, 2011). In this study, we use isoform 2 of HGSNAT, which is produced as a 635 amino acid protein containing a 30 amino acid N-terminal signal peptide, a ∼110 amino acid long luminal domain, and 11 transmembrane helices (TMs). HGSNAT is believed to be expressed in the cell as an immature precursor protein that localizes as an *N*-glycosylated dimer on the lysosomal membrane (Durand *et al,* 2010; Fan *et al,* 2011). The localization of HGSNAT happens via the adaptor protein-mediated pathway, aided by the lysosomal targeting motifs *[DE]-XXXL[LI]* (^204^ETDRLI^209^) and *YXXØ* (^624^YILYRKK^630^) present towards the N- and C-terminus of HGSNAT respectively (Rudnik & Damme, 2021; Schwake *et al*, 2013). Deleting the C-terminal lysosomal sorting signal in HGSNAT retains the protein in the plasma membrane (Bonifacino & Traub, 2003; Durand *et al,* 2010). Once targeted to the lysosomal membrane, HGSNAT, like a few other lysosomal membrane proteins, is proteolyzed by unidentified acid proteases in the lysosomal lumen into two unequal fragments - a smaller luminal N-terminal α-HGSNAT and a larger transmembrane C-terminal β-HGSNAT that continue to co-localize despite the proteolysis (Durand *et al,* 2010; Fan *et al,* 2011; Rudnik & Damme, 2021; Steenhuis *et al*, 2012). Upon proteolytic maturation, it is believed that HGSNAT is assembled as a hetero oligomer of α-& β-HGSNAT chains (Durand *et al,* 2010; Feldhammer *et al*, 2009b). However, the essentiality of proteolytic cleavage and oligomerization for HGSNAT activity remains debated as endoplasmic reticulum (ER) retained monomeric HGSNAT was also shown to be active (Durand *et al,* 2010; Fan *et al,* 2011).

So far, over 70 unique mutations in the *HGSNAT (TMEM76)* gene have been identified. These mutations span the entire sequence and include deletions, nonsense mutations, splice-site variants, and silent and missense mutations (Canals *et al*, 2011; Fan *et al,* 2006; Fedele & Hopwood, 2010; Feldhammer *et al*, 2009a; Feldhammer *et al,* 2009b; Hrebicek *et al,* 2006; Huizing & Gahl, 2020). The majority of the mutations that cause MPS IIIC are missense mutations that result in HGSNAT folding and localization defects, making them ideal targets for pharmacochaperone therapy. Site-specific inhibitors of HGSNAT activity have been explored as agents to rescue the misfolded conformation of certain missense variants, thereby restoring the partial *N*-acetyltransferase activity (Coutinho *et al,* 2012; Fedele & Hopwood, 2010; Feldhammer *et al,* 2009b; Pan *et al*, 2022). Despite its clinical relevance, the structure of HGSNAT and the mechanism of *N*-acetyltransferase activity are poorly understood. Here, using single-particle cryo-electron microscopy (cryo-EM), we report the high-resolution structure of full-length HGSNAT in a complex with acetyl-CoA. This is the first structure of a member of the transmembrane acyl transferase (TmAT) superfamily (classes 9.B.97 and 9.B.169 in the Transporter Classification Database (TCDB)) (Saier *et al*, 2021). The HGSNAT-acetyl-CoA complex structure presented here provides a high-resolution snapshot of the first step in HGSNAT catalyzed acetyltransferase reaction, and reveals critical cofactor binding amino acids in the HGSNAT active site. In addition, our structure also provides molecular insights into the impact of MPS IIIC-causing mutations on acetyl-CoA binding and the overall architecture of the protein.

## Results

### Purification of dimeric HGSNAT

Two groups have previously reported purification of HGSNAT in varying oligomeric forms (Durand *et al,* 2010; Fan *et al,* 2011; Feldhammer *et al,* 2009b). The first group used non-ionic surfactants, NP-40 and Triton X-100, in their purification experiments and observed dimers and hexamers of HGSNAT. The PEG-based headgroup in these detergents is relatively bulky and yields micelles of variable sizes (45-100 kDa) and has been known to cause sample heterogeneity in eukaryotic membrane proteins (Orwick-Rydmark *et al*, 2016). The second group purified monomeric HGSNAT using non-ionic detergent DDM, which forms a relatively uniform, albeit large, micelle (98kDa). However, the purification process involved two overnight incubation steps – the first in 1% and the second in 0.2% DDM. Both groups used transient transfection to express the protein in COS-7 or HeLa cells, respectively. All these factors could impact the monodispersity of purified HGSNAT. To identify the ideal conditions for expression and purification of monodisperse HGSNAT, we transfected HEK293 GnTI^-^ cells with plasmids expressing either N-terminal or C-terminal GFP-fusion of HGSNAT and monitored GFP fluorescence in detergent-solubilized HGSNAT lysates by fluorescence-detection size-exclusion chromatography (FSEC) (Kawate & Gouaux, 2006). We noticed that the position of GFP did not alter the expression of HGSNAT (Fig S1A). However, the HGSNAT dimer model predicted using ColabFold suggested that the C-termini of the protomers lie at the dimer interface (Mirdita *et al*, 2022). Thus, for large scale production of HGSNAT for structural studies, we expressed N-terminal StrepII-tag-GFP-HGSNAT fusion. Large quantities of GFP-HGSNAT fusion were produced using baculovirus mediated transduction of HEK293 GnTI^-^ cells (Goehring *et al*, 2014). We found that reducing the temperature of the suspension cell culture to 32°C at about 8-10 hr after transduction, or transient transfection, resulted in a better yield than continuing to grow cells at 37°C (Fig S1B). To identify the ideal conditions for the purification of HGSNAT, we screened various detergents. In most of the detergent conditions we tested, HGSNAT was predominantly dimeric (Fig S1C-H). Then, we performed a single-point thermal melt test of solubilized cell lysates to compare the relative stability of HGSNAT in different detergents. The rationale being that heating should exacerbate the instability of the protein in a said condition. HGSNAT appeared most stable in digitonin (Fig S1I-L). Based on our FSEC analysis of relative thermal stability and homogeneity, we purified HGSNAT in digitonin. We observed a dimer in our chromatography experiments and a band corresponding to a monomer on SDS-PAGE (Fig S1M-O).

### Architecture of HGSNAT

We solved the structure of HGSNAT in complex with acetyl-CoA (HGSNAT-ACO complex) to a global resolution of 3.26 Å, with the transmembrane domain (TMD) being better resolved than the luminal domain (LD) (Fig S2 and Fig S3A-C). This is the first experimentally determined structure of a member of the transmembrane acetyltransferase protein family (TmAT). Our structure reveals that HGSNAT is a dimer where the protomers are related to each other by 2-fold rotational symmetry, and the C2 axis of rotation is perpendicular to the plane of the membrane (Figs 1A-1C). Each polypeptide of HGSNAT has an N-terminal LD followed by 11 TMs that comprise the TMD, which is ensconced in the lysosomal membrane. We could model all the secondary structure elements of the protein unambiguously except the first 48 amino acids including the N-terminal signal peptide and the cytoplasmic loop 1 (CL1; L183-L236). The regions that are poorly resolved are the β-turn that connects π7 and π8 (N140-E148) the C-terminal half of TM1 (V176-F181) (Fig 1A, Fig S3D, and Fig S3E). The C-terminus of the protein lies at the dimer interface but is unlikely to be directly involved in dimerization as the C-termini of the protomers lie ∼25 Å away from each other, pointing towards the central acetyl-CoA binding site (ACOS) (Fig 1B and Fig 1C).

**Figure 1:**
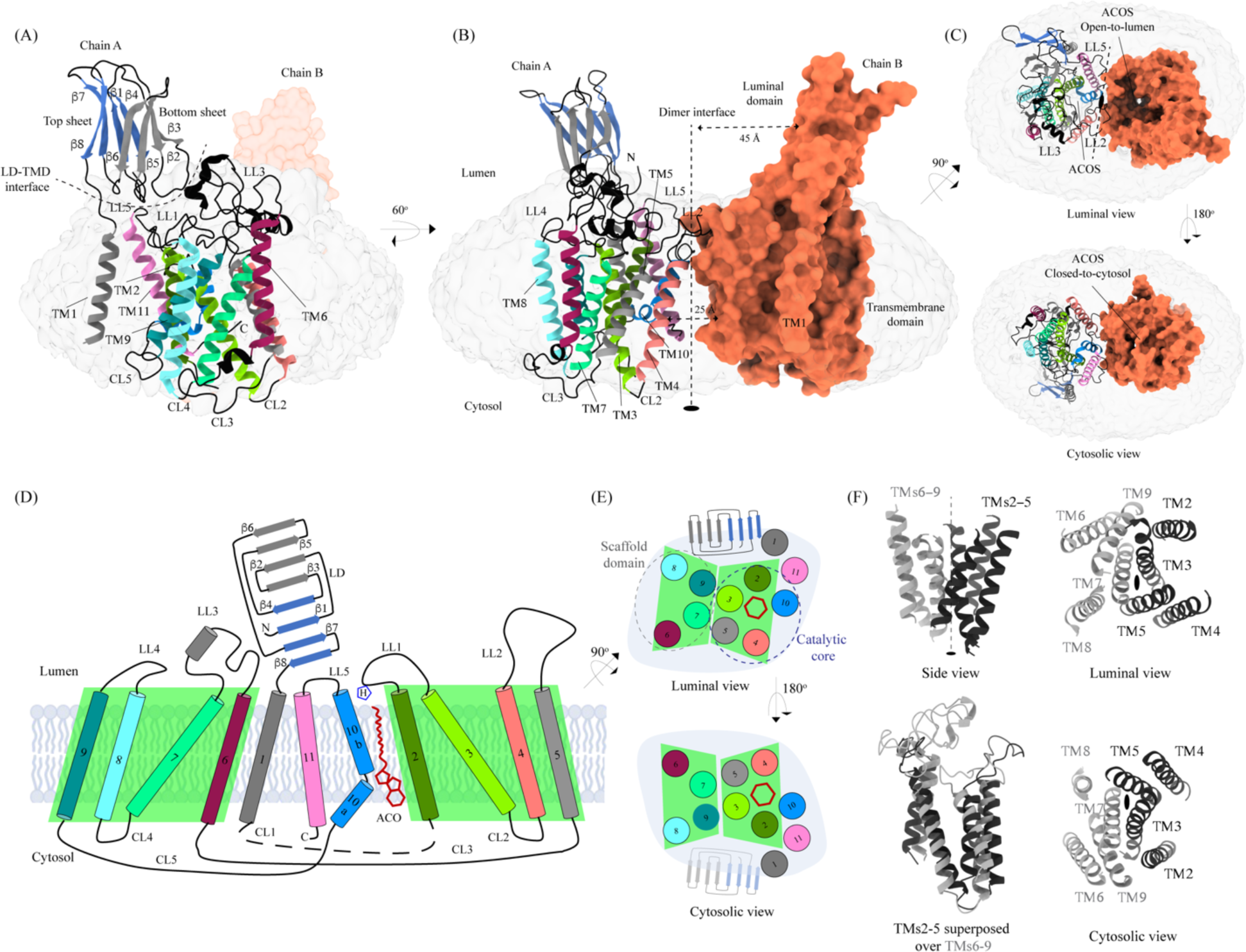
Structure of HGSNAT. Panels **(A)** and **(B)** show two different orientations of HGSNAT dimer that highlight (dashed lines) the LD-TMD interface and dimer interface respectively. Micelle is displayed in gray. Chain A is displayed as a cartoon and chain B as orange surface. All the luminal loops (LLs), cytosolic loops (CLs), and the loops that connect β-sheets are shown in black. The top and bottom sheets in the luminal domain (LD) are colored blue and gray, respectively. The two-fold rotation axis is displayed as a dashed line with an ellipsoid. **(C)** Luminal (top) and cytosolic (bottom) views of the protein. The surface representation of chain B suggests that the acetyl-CoA binding site (ACOS) is more accessible from the luminal side (top) than the cytosolic side (bottom). **(D)** 2D topology of HGSNAT and YeiB family. The helices and strands in the topology are colored similarly to the 3D structure. TMs 2-5 and 6-9 form two bundles (4+4), highlighted by green parallelograms, that are related to each other by a 2-fold rotation parallel to the plane of the membrane. TMs 1, 10, and 11 do not seem involved in this internal symmetry, with TM10 being bent in the plane of the membrane into two halves TM10a and TM10b. The relative position of bound ACO and active site H269 of LL1 are indicated. **(E)** Luminal (top) and cytosolic (bottom) views of the protein topology. TMs 2-5 and TM10 enclose ACOS (red hexagon) and are referred to as catalytic core (blue dashed oval). TMs 6-9 will be referred to as scaffold domain (gray dashed oval). **(F)** 4+4 bundle formed by TMs 2-5 (black) and TMs 6-9 (gray) are related by a 2-fold rotation. The last sub-panel (bottom left) shows a superposition of TMs 2-5 on TMs 6-9.

**Table 1:** Cryo-EM data collection, processing, and validation statistics.

|  | EMDB-41620 and PDB-8TU9 |
| --- | --- |
| ***Data collection and processing*** |  |
| Magnification | 105,000x |
| Voltage (kV) | 300 |
| Data collection mode | Super-resolution |
| Electron exposure (e-/Å<sup>2</sup>) | 50 |
| Defocus range (μm) | -1.0 to -2.5 |
| Physical Pixel size (Å) | 0.848 |
| Symmetry imposed | C2 |
| Initial particle images (no.) | 3,325,732 |
| Final particle images (no.) | 57739 |
| Map resolution (unmasked, Å) at FSC 0.143 | 3.7 |
| Map resolution (masked, Å) at FSC 0.143 | 3.3 |
| Map resolution range (Local resolution) | 2.5-4.5 |
| ***Refinement*** |  |
| Map sharpening B factor (Å<sup>2</sup>) | -30 |
| **Model composition** |  |
| *Chains* | 4 |
| *Atoms* | 8284 (Hydrogens: 0) |
| *Residues* | Protein: 1066 Nucleotide: 0 |
| *Water* | 0 |
| *Ligands* | ACO: 2 |
| **Bonds (RMSD)** |  |
| *Length (Å) (# > 4sigma)* | 0.008 (0) |
| *Angles (°) (# > 4sigma)* | 1.480 (28) |
| MolProbity score | 1.25 |
| Clash score | 2.32 |
| **Ramachandran plot (%)** |  |
| *Outliers* | 0.00 |
| *Allowed* | 3.59 |
| *Favored* | 96.41 |
| **Rama-Z (Ramachandran plot Z-score, RMSD)** |  |
| *whole (N = 1058)* | -0.40 (0.24) |
| *helix (N = 486)* | 1.03 (0.21) |
| *sheet (N = 114)* | 1.69 (0.51) |
| *loop (N = 458)* | -2.35 (0.23) |
| Rotamer outliers (%) | 0.00 |
| Cβ outliers (%) | 0.00 |
| **Peptide plane (%)** |  |
| *Cis proline/general* | 0.0/0.0 |
| *Twisted proline/general* | 0.0/0.0 |
| Cα BLAM outliers (%) | 4.38 |
| ADP (B-factors) |  |
| *Iso/Aniso (#)* | 8284/0 |
| *Protein (min/max/mean)* | 10.47/125.39/28.58 |
| *Ligand (min/max/mean)* | 16.71/16.71/16.71 |
| ***Model vs. Data*** |  |
| CC (mask) | 0.78 |
| CC (box) | 0.63 |
| CC (peaks) | 0.58 |
| CC (volume) | 0.72 |
| Mean CC for ligands | 0.77 |

#### Transmembrane domain (TMD)

The TMD of HGSNAT protomer comprises of a central 4+4-fold, where the TMs 2-5 are related to TMs 6-9 by a 2-fold rotational pseudo-symmetry with the axis of rotation being perpendicular to the plane of the membrane (Figs 1D-F). TMs 2-5, along with TM10, form a ‘catalytic core’ and enclose the ACOS accessible via the cytosol and lumen along the dimer interface. ACOSs of the protomers lie on either side of the dimer interface axis. TMs 6-9, along with TM11, form a ‘scaffold domain’ separated from the dimer interface by the catalytic core (Fig 1E). The third luminal loop (LL3) connecting TM6 and TM7 forms a lid that limits the access to the cavity between the catalytic core and the scaffold domain from the luminal side (Figs 1A, 1C, and 1D). The TMD region comprising TM2-TM11 of HGSNAT is predicted to be evolutionarily conserved across HGSNATs from other kingdoms. Owing to the unique architecture we have named this novel fold as the transmembrane N-acetyltransferase (TNAT) fold (Fig S4A and Fig 4A). Although the resolution for the C-terminal half of TM1 is poor, it is sufficient to position the helix separately from the rest of the TMD (Fig 1A). TM1 is connected to TM2 by ∼50 amino acid long CL1 (Fig 1D). LL1-LL3 and LL5 form the boundaries for the luminal entrance of ACOS. The C-terminus, CL2, and a part of CL1 form the boundaries of the cytosolic entrance of ACOS. The ACOS in our structure is relatively more accessible from the luminal side than the cytosolic side, as the nucleoside head group of the bound acetyl-CoA blocks the cytosolic entrance of ACOS (Fig 1C and Fig S5). Thus, the HGSNAT-ACO complex is in a conformation where the active site is readily accessible for the binding of the second substrate from the lysosomal lumen.

#### Luminal domain (LD)

The LDs of the protomers lie diagonally opposite to each other ∼45 Å away from the dimer interface and are not involved in dimerization (Fig 1B). Located between the N-terminal signal peptide and the TM1, LD is a ∼110 amino acid long β-sandwich made of two beta sheets of 4 strands each where strands β1, β4, β7, & β8 form the mixed β-sheet on top (Figs 1A-E, blue), and the strands β2, β3, β5, & β6 arrange as bottom anti-parallel β-sheet (Figs 1A-E, gray). The mixed β-sheet is arranged such that the order of the strands is β4-β1-β7-β8, with β1 & β7 being parallel. β8 of LD is connected to TM1 (Fig 1B, 1D, and 2A). LD has two predicted disulfide bonds – one in the β-turn that connects β2 and β3 (C76-C79), and the other between the strands β6 and β8 (C123-C151) holding the two sheets together. The resolution of the LD domain in our structure allows us to model only one (C76-C79) of these unambiguously (Fig 2B). HGSNAT is produced as a pro-protein that is believed to get proteolyzed into two fragments, HGSNAT-α and HGSNAT-β, of unequal sizes, which remain together (Durand *et al,* 2010; Fan *et al,* 2011). It is unclear if HGSNAT-α is made of just LD or LD and TM1, and if HGSNAT-β is made of all TMs or only TMs2-11 (Fig 2A and Fig S4). While our structure has relatively poor local resolution at the predicted protease sites, making it difficult to map them unambiguously, the FSEC nor the SDS-PAGE analyses of the purified protein indicate that the recombinant HGSNAT is not proteolyzed (Fig S1). We believe the relatively low local resolution of LD is because of the flexibility introduced by GFP fused to the N-term of HGSNAT.

**Figure 2:**
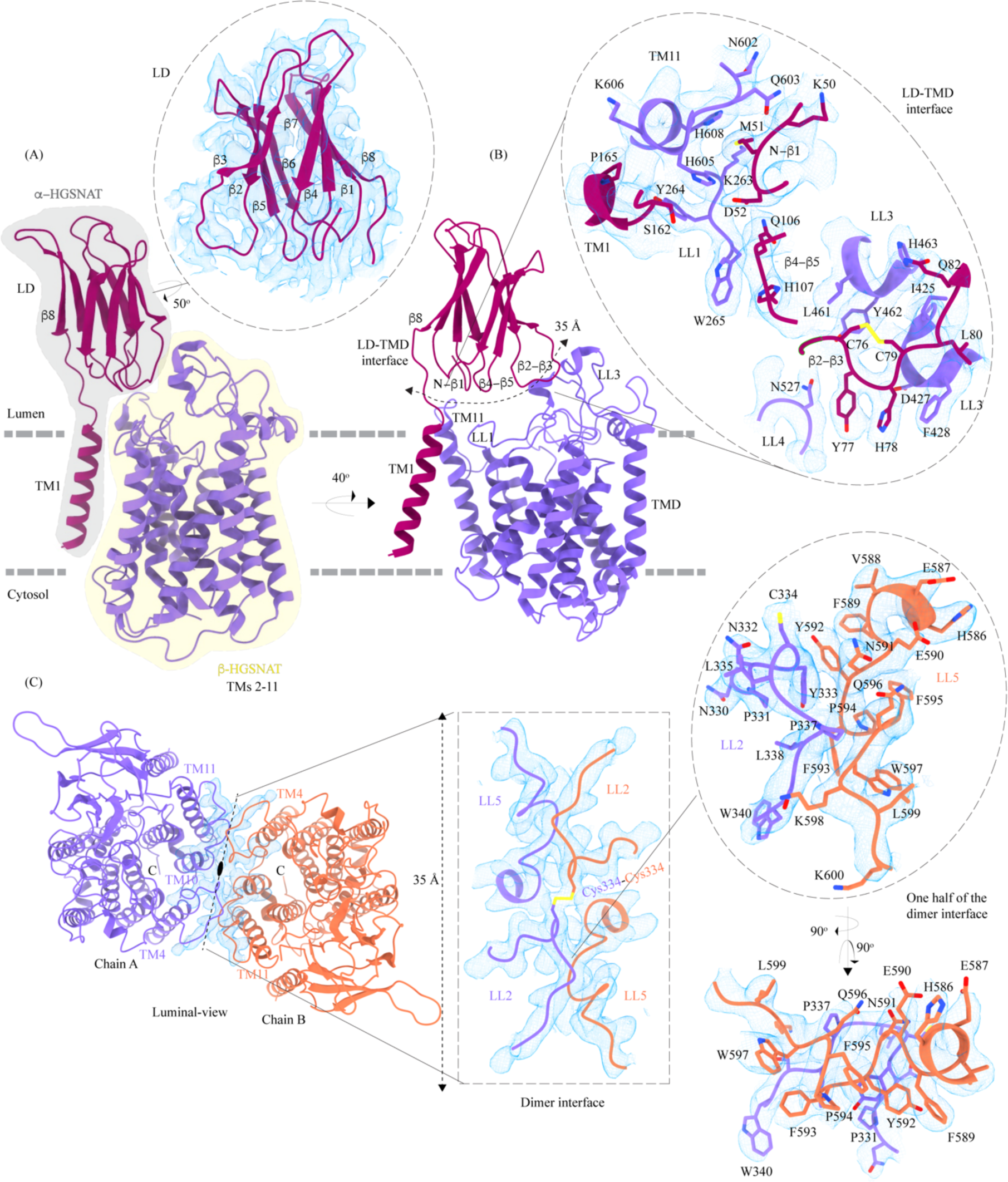
Domain organization, and LD-TMD and dimer interfaces of HGSNAT. **(A)** HGSNAT is predicted to be proteolyzed into two chains of unequal size - α-HGSNAT (dark magenta cartoon, gray shaded area) and β-HGSNAT (purple cartoon, yellow shaded area). The site for proteolysis remains debated. Based on our structure and prediction of HGSNAT structures from other kingdoms (Fig S4), we have represented α- and β-HGSNAT fragments as shown in panel A. The inset (dashed oval) shows the luminal domain (dark magenta) fit to cryo-EM density (blue; display level 0.21 of the composite map in ChimeraX) (Fig S3). The lysosomal membrane is shown as a dashed gray line. **(B)** LD-TMD interface is highlighted (dashed line). Inset highlights the residues that interact at the LD-TMD interface, and cryo-EM density for the same (blue; display level 0.25 of the 3.26 Å C2 refined map in ChimeraX). C76-C79 disulfide of β2-β3 turn is shown as yellow sticks, while the residue sidechains are colored the same as their secondary structure elements, with heteroatoms highlighted. **(C)** Luminal-view of the protein with dimer interface highlighted (dashed line). Inset (dashed rectangle) highlights LL2 and LL5 that line the dimer interface, and the C334-C334 inter-chain disulfide (yellow) between the chains A (purple) and B (orange). The dashed oval inset shows one-half of the dimer interface with LL2 and LL5 of chains A and B, respectively, contributing other hydrophobic interactions that stabilize the dimer interface. The cryo-EM density in panel C is displayed as blue mesh (display level 0.22 of the C2 refine map in ChimeraX).

#### LD-TMD interface

LD interacts with TMD at multiple sites via an extensive interaction network that spans ∼35 Å, comprised of a salt bridge (D52-H605), hydrogen bonds, and dipole-dipole and hydrophobic interactions (Fig 2B and Fig S6A). The N-terminus of TM11 interacts with the N-terminus of β1 and LL1. LL1 also interacts with the N-terminus of TM1 and the β4-β5 turn. The β2-β3 turn, including the C76-C79 disulfide, interacts extensively with LL3 and a part of LL4. The β2-β3 turn-LL3-LL4 interaction is also stabilized by a 3π-network formed by the stacking of Y77, H78, & F428 side chains. The hydrogen bond network that stabilizes the LD-TMD interaction is formed between the amino acids M51, H78, and S162 of LD with H605 and Q603, D427 and F428, and Y264 of the TMD respectively (Figs 2B and S6A). The LD-TMD interface is separated from the dimer interface by the central catalytic core (Figs 2B and 2C).

#### Dimer interface

The dimer interface is spread across ∼ 35 Å towards the luminal side of the protein, perpendicular to the C2 rotation axis (Fig 2C). Although the TMs 4, 10, and 11 from both protomers lie on either side of the dimer interface, they do not directly interact to stabilize the dimer interface. The primary mediator of dimerization is the disulfide between C334s within the LL2s of each protomer (Fig 2C, rectangle inset). In addition to the C334-C334 disulfide, the dimer interface is also stabilized by an extensive ρχ-ρχ interaction network formed between the aromatic residues of LL2 (332-340) of one protomer and the aromatic residues of LL5 (588-598) of the other protomer (Fig 2C and Fig S6B). Notably, the aromatic bulky side chains at the interface are buried in the membrane, and the hydrophilic residues face the lumen (Fig 2C, oval inset). Y333, L338, and S339 of LL2 from one protomer form a series of hydrogen bonds with F593 and K598 of LL5 of the other protomer, which adds to the stabilization of the dimer interface (Figs 2C and S6B). In all the detergents we tested, HGSNAT eluted as a dimer, a testimony to the extensive side-chain interaction network. The 2-fold rotational symmetry between the protomers juxtaposes the ACOSs of protomers connecting them with each other on the cytosolic side (Figs 1 and 2C). This interconnected space is partitioned by lipids, preventing the diffusion of ligands from one active site to the other within a dimer (Fig S7). We generated two HGSNAT mutants of residues at the dimer interface – C334A and F593A. C334 from each protomer are linked in a disulfide bond, and F593 forms hydrogen bonds with multiple residues from the opposite protomer (Figs 2C and S6B). In both these mutants, the overall expression of HGSNAT was not reduced. In case of F593A, the stability of dimer also remained unaffected (Figs S8A, S8H, S8I, and S8N). While both these mutants still expressed predominantly as dimers even in the presence of 1% digitonin, the C334A mutant eluted as a monomer upon heat treatment (Figs S8E, S8H, S8I, and S8K). As C334A breaks a covalent disulfide link between the protomers, in the absence of this disulfide, the extensive side-chain interaction network stabilizes the dimer interface. Heating destabilizes these interactions, resulting in a monomeric C334A.

### Acetyl-CoA binding site (ACOS)

We used MOLEonline to predict the presence of a ∼75Å pore within the TMD, that begins at the cytosolic side and extends to the luminal side (Pravda *et al*, 2018) (Fig S5A). The pore predicted by MOLEonline is lined by highly conserved residues of the TMs 2-5 and TM 10 (Fig 3 and Fig S5C). We note that the helix TM10 is bent by ∼80° such that the luminal-side 2/3^rd^ (TM10b) of the helix is parallel to TM2 and the cytosolic-side 1/3^rd^ (TM10a) of the helix protrudes between TM2 and TM11 (Fig 3A). The bending of TM10 is aided by P575 and G576, residues known to induce helix breaks and kinks (Javadpour *et al*, 1999; Ulmschneider & Sansom, 2001). We find a density in our cryo-EM map overlapping the MOLEonline prediction in both protomers, spanning the entire length of TMD, that allowed us to unambiguously model a single acetyl-CoA, such that the 3’, 5’-ADP nucleoside head group is towards the cytosol and the acetyl group is hydrogen bonded (∼3.5Å) to N258 of TM2 that is located at the luminal entrance of ACOS. On the cytosolic side, the orientation of TM10a between TM2 and TM11 seemed to have allowed the positioning of the 3’, 5’-ADP nucleoside head group in the space created by bending away of TM10a from the central axis of the catalytic core. The catalytic H269, which is predicted to be acetylated during the acetyltransferase reaction, is ∼4.5 Å away from the acetyl group of acetyl-CoA, nestled within the negative charge on the luminal opening, and is ergonomically positioned to perform acetyltransferase reaction in the presence of the acetyl group acceptor (Fig 3A). The binding pocket, like the rest of the protein, shows a polarity of charge. As we move from the cytosolic side to the luminal side, the charge changes from positive to neutral to negative (Figs 3B, S5A, D). The cytosolic opening of the site is comprised of basic and nonpolar amino acids, making it an ideal pocket for binding the 3’, 5’-ADP head group. The pantothenate group of acetyl-CoA is supported by a network of nonpolar amino acids at the center of the binding pocket (Figs S5A, D). A series of conserved salt bridges stabilize the cytosolic and luminal entrances of the pore. The luminal entrance is lined with salt bridges between the residues H269, D279, R344, D469, E471, H586, and E587. The cytosolic entrance is lined with salt bridges between the residues R239, D244, R247, R317, E363, K491, and K634 (Figs S5C, D). A portion of CL1 towards the TM2 and C-terminus of the protein also seems to be a part of the cytosolic entrance of the ACOS.

**Figure 3:**
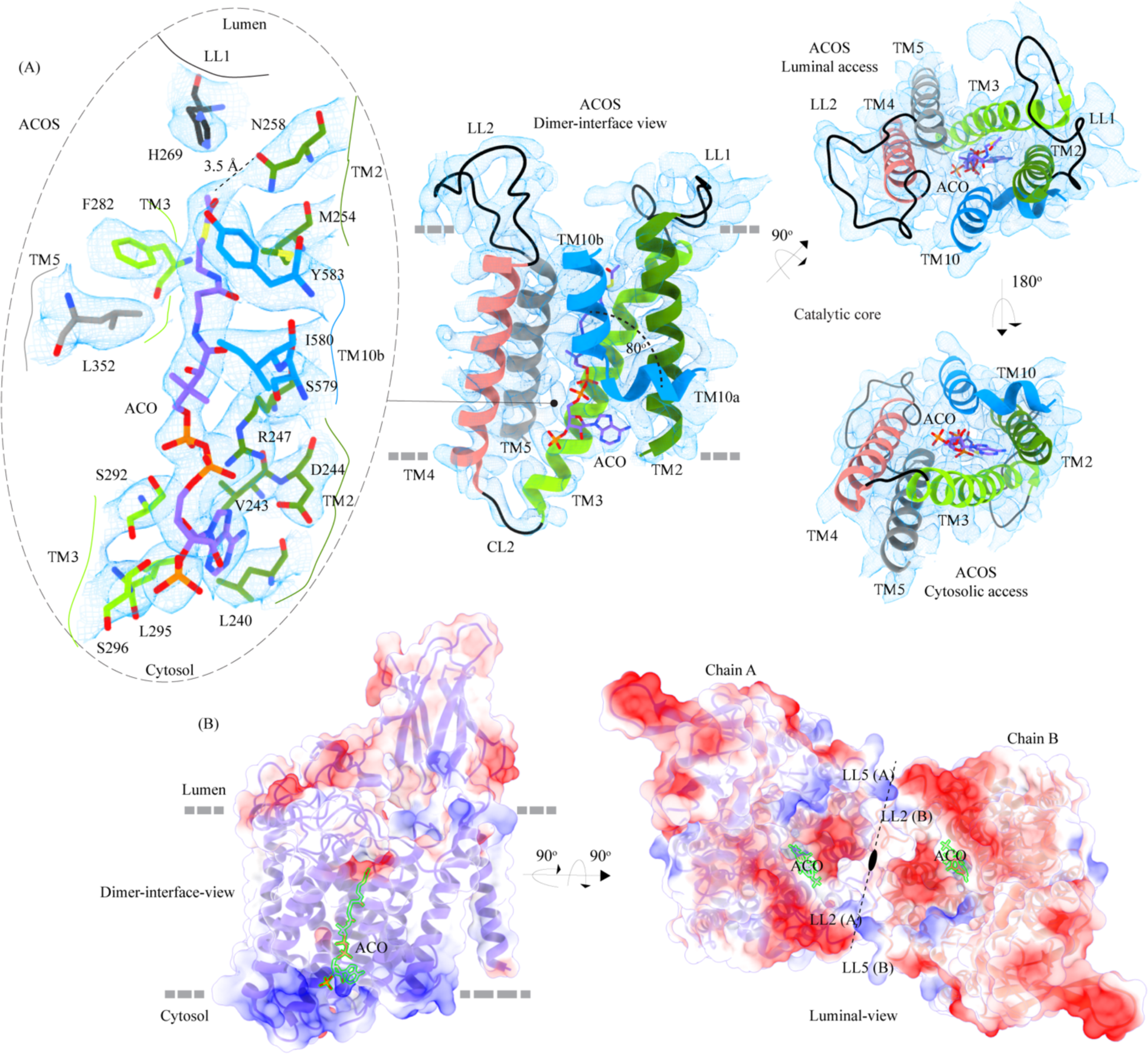
Acetyl-CoA binding site (ACOS) **(A)** Catalytic core (chain A) of HGSNAT comprised of TMs 2-5 and TM 10. LLs and CLs are shown in black, and the helices are colored as in Fig 1. Acetyl-CoA (ACO) is colored (purple), the same as chain A in Fig 2 with heteroatoms highlighted. The inset (dashed oval) shows ACOS and highlights the amino acids of HGSNAT that interact with ACO. The amino acids are colored same as the corresponding TMs, with heteroatoms highlighted. Cryo-EM density for ACOS is displayed as blue mesh (display level 0.3 of the 3.26 Å C2 refine map in ChimeraX). ACO could be modeled into the densities at chain A and B ACOSs with a mean correlation coefficient (CC) of 0.77. The nucleoside headgroup of ACO plugs in the cytosolic access of ACOS, and the luminal access seems relatively more accessible. **(B)** Electrostatic potential and surface charge distribution of HGSNAT, with the surface display colored based on the potential contoured from -10 kT (red) to +10 kT (blue). ACO bound at the ACOS is highlighted in golden yellow. Luminal and cytosolic sides of the protein show a conspicuous polarity. The lysosomal membrane is shown as a dashed gray line in both sub-panels.

### Conservation and homology

Sequence conservation between HGSNAT and other known acetyl-CoA binding proteins or transferases is poor. There are no structural homologs of HGSNAT, and a search within the database of known structures using the HGSNAT TMD by the Dali server resulted in hits with sequence identity of <17% across alignments of <25% sequence length (Holm *et al*, 2023). Even the search with just the luminal domain, which appeared to be a classic two-sheet β-sandwich, yields hits with <15% identity over alignments of at least 70% sequence lengths (Table S1, Fig S4 C, D). We used ModelAngelo, a machine-learning based *de novo* automatic model building algorithm to build initial HGSNAT model into the cryo-EM density (Jamali *et al*, 2023). The model built by ModelAngelo into the experimental data superposed well with the HGSNAT model as predicted by AlphaFold with a Cα RMSD of ∼1.7 Å over 533 amino acids (Fig S4A) (Jumper *et al*, 2021; Varadi *et al*, 2022). However, unlike the AlphaFold model of Isoform 1, our expression construct lacks the extended signal peptide (N-terminal 28 amino acids). In addition, we do not observe any density to model CL1 that connects TM1 and TM2. HGSNATs of representatives from archaea, bacteria, and plants suggest that LD and TM1 are absent in other kingdoms. These homologs superpose onto the TMD part of HGSNAT, especially TM2-11 with Cα RMSDs of 1-1.4 Å over 350 amino acids (Fig S4). TmAT superfamily of membrane proteins consists of two subfamilies of integral membrane proteins consisting of 8-12 TMs – the acyltransferase-3/acetyl-CoA transporter (ATAT) family (TCDB: 9.B.97) and the integral membrane acetyltransferase (YeiB) family (TCDB: 9.B.169). HGSNAT belongs to the YeiB family. A meaningful sequence alignment could not be generated between the members of these two families. ATAT family of transporters seem to have a single extra-membranous domain, like HGSNAT (Fig S4B). However, based on AlphaFold prediction, this extra-membranous domain seems to be a α-β-α sandwich with a single 5-stranded β-sheet sandwiched between two helical domains. Incidentally, a bent helix near the predicted acetyl-CoA binding site is also observed in the ATAT family, suggesting that this could be a conserved feature amongst the members of TmAT superfamily (Newman *et al*, 2023). We used ConSurf to estimate sequence conservation in HGSNAT based on an automatic sequence alignment algorithm (Ashkenazy *et al*, 2016). It appears that the LD and TM1 regions have poor sequence conservation compared to the rest of the protein (Fig 4A).

**Figure 4:**
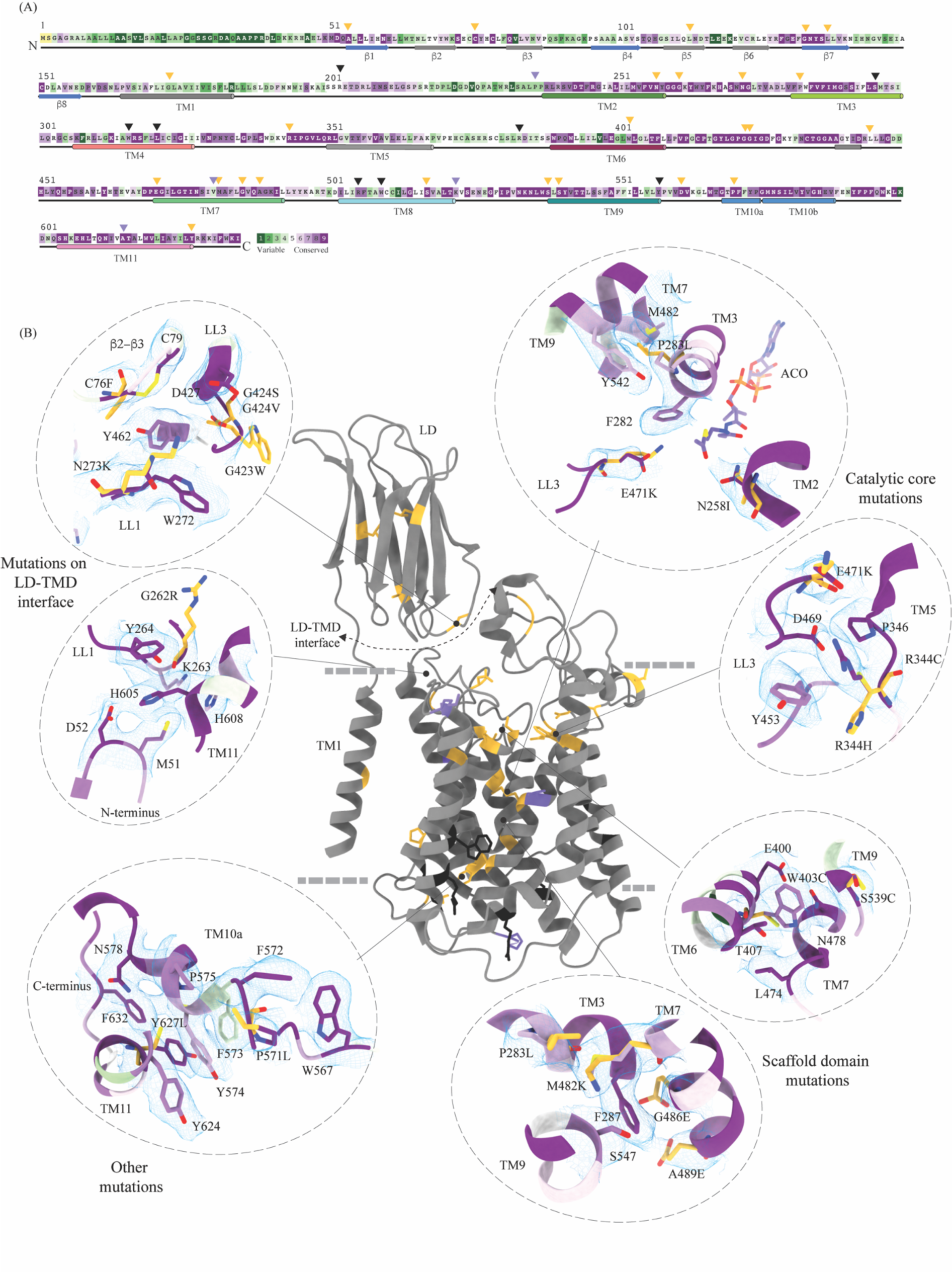
Molecular basis for MPS IIIC mutation-induced dysfunction. **(A)** Evolutionary sequence conservation of HGSNAT. Amino acids are color coded according to the conservation scores generated by ConSurf webserver using a Clustal multiple sequence alignment of homologs identified by PSI-BLAST (Ashkenazy *et al,* 2016). The positions of the mutations - missense (orange), nonsense (black), and polymorphisms (purple) – are indicated on the sequence by triangles. **(B)** MPS IIIC-causing mutations mapped on the HGSNAT structure. The color coding of the positions is the same as in panel A. Some of the missense mutants are highlighted in the insets (dashed ovals). We grouped them based on their position within the protein – LD-TMD interface, catalytic core, scaffold domain, and other C-terminal mutations. The insets show the 3D environment of the mutant sites on the wild-type HGSNAT color coded as per their evolutionary sequence conservation scores, and the potential disturbance to it caused by the mutation (orange side chains). The coordinates for mutant side chains were generated based on wild-type HGSNAT structure as input in FoldX webserver (Schymkowitz *et al,* 2005).

### The basis for mutation-induced dysfunction and destabilization of HGSNAT

We mapped the known clinical mutations (missense, nonsense, and polymorphisms) of HGSNAT onto its three-dimensional structure (Canals *et al,* 2011; Fan *et al,* 2006; Fedele & Hopwood, 2010; Feldhammer *et al,* 2009a; Feldhammer *et al,* 2009b; Hrebicek *et al,* 2006; Huizing & Gahl, 2020). Almost all the mutations fall within the conserved regions on HGSNAT (Fig 4A). The nonsense mutations seem to be located more on the cytosolic side, and the missense mutations seem to be populated on the luminal side of HGSNAT (Fig 4B). We estimated the extent of destabilization or stabilization introduced by these variants on HGSNAT structure by FoldX, a force field algorithm to evaluate the effect of mutations on the stability and dynamics of proteins (Table S2) (Schymkowitz *et al*, 2005). We classified these mutations into four groups based on their position on the structure and found that most destabilizing mutations appear to be concentrated near the LD-TMD interface (Fig 4 and Table S2). Our structure provides the basis for destabilization or dysfunction induced by missense mutations in HGSNAT.

#### Catalytic core mutations

Five missense mutations - E471K (LL3), R344C/R344H (LL2), P283L (TM3), and N258I (TM2) - have been identified in the residues that line the ACOS (Figs 4B, 3A, and S5C). E471 is close to the acetyl group of acetyl-CoA, and acidic to basic side chain substitution in an E471K mutation could impact the binding affinity of acetyl-CoA. N258 side chain can act as both hydrogen acceptor or donor and is hydrogen bonded to acetyl-CoA in the structure. N258I mutation could directly impact the binding of acetyl-CoA and acetyltransferase activity. R344 forms salt bridges with E469 and E471 that stabilize the luminal entrance of ACOS. The mutations of R344C/R344H could affect the integrity of the luminal entrance of ACOS. In addition to being involved in acetyl-CoA binding, all these three positions – E471, R344, and N258 – are also highly conserved (Fig 4A, B). P283 is not directly involved in the binding of acetyl-CoA. However, the residues flanking P283 – V281, F282, F285, I288, and M289 – all interact with the pantothenate group of acetyl-CoA (Fig 3A and Fig S5D). Proline residues define helix conformation, and mutation of a relatively conserved proline to an aliphatic leucine could alter the TM3 conformation on the luminal side and thus affect acetyl-CoA binding. FoldX prediction indicates that P283L is the most destabilizing of all ACOS mutations perhaps because it impacts the TM3 conformation. In contrast, other catalytic core mutants only show a potential to impact the binding of acetyl-CoA. We generated N258I and R344H mutants to test the effect of these substitutions on the expression and stability of HGSNAT. We noticed that these substitutions did not reduce the overall expression of HGSNAT (Fig S8A). R344H mutant, upon solubilization in 1% digitonin, showed slightly enhanced aggregation but the overall stability of R344H mutant was not altered drastically, compared to WT HGSNAT (Figs S8F, S8I, and S8L). Although N258I mutant showed expression comparable to WT HGSNAT, as evident by total fluorescence measurements of solubilized cell lysates, we could not observe any peak for N258I nor free GFP in FSEC (Figs S8A and S8C). We hypothesize that N258I substitution directly affected the substrate binding and stability. As a result, upon solubilization followed by ultracentrifugation, the unstable protein got pelleted out of the solution (Figs 4B and S8C).

#### Mutations at the LD-TMD interface

C76F, G262R, N273K, G423W, and G424S/G424V are all missense mutations that are on the LD-TMD interface, and our FoldX based analysis indicates these mutations to be the most destabilizing among the ones that we listed. N273K is predicted to be least destabilizing by FoldX, as all these mutations, except N273K, result in charge reversal and drastic change in the side chain size, leading to steric clashes and breakdown of existing interactions (Fig 4B and Fig 2B). Although none of these missense positions at the LD-TMD interface, except C76, are directly involved in LD-TMD interface contacts, the drastic side chain changes in the vicinity of the residues directly involved are expected to destabilize the interaction (Fig 2 and Fig S6). For example, the glycine residues (G262, G423, and G424) lie within pockets lined by aromatic side chain containing amino acids, and the substitution of such a residue with a bulky side chain will cause a steric clash, destabilizing those pockets of interaction (Fig 4B). C76F mutation reduces the expression of the protein, suggesting that LD-TMD interaction is essential for proper folding and stability of HGSNAT (Figs S8A and S8B).

#### Scaffold domain mutations

W403C, M482K, G486E, A489E, S518F, S539C, and S541L are all mutations that occur in the scaffold domain (TM6-TM9) of HGSNAT. FoldX predicts these mutations to be only mildly destabilizing. For example, the change in the charge and size of S539C or W403C mutation is not drastic enough to destabilize TM6 or TM9 helices. Even in the cases where there are large substitutions, for example, G486E or A489E, the substituted side chains face away from the core of the protein and are involved in minimal interactions or clashes (Fig 4B). Corroborating the FoldX prediction, we noticed that W403C was mildly destabilizing. While the overall expression of the W403C mutant was not affected, the thermal stability was reduced compared to WT HGSNAT (Fig S8A, S8G, S8I, and S8M).

#### Other mutations

Mutations in the LD domain, such as L113P, G133A, and L137P, seem to face the lumen and are involved in minimal interactions. Amongst the mutations that we analyzed by FoldX, none of them were on the dimer interface. However, two mutations, P571L and Y627C, towards the cytosolic ends of TM10 and TM11 seem closer to the dimer interface on the cytosolic side. However, they are not directly involved in dimerization. These ring side chain amino acids are part of an elaborate ε-ε-network stabilizing TM10a and the C-terminus in positions that make space for the nucleoside head group at the ACOS (Fig 4B and Fig 3A). Drastic mutations in this region could impact acetyl-CoA binding and thereby the conformational stability of catalytic core.

### Mechanism of acetyltransferase reaction

HGSNAT, in this report, was purified at pH 7.5. During the purification process, we did not add acetyl-CoA to the buffer. However, we were able to confirm the presence of endogenously bound acetyl-CoA in our cryo-EM sample by LC-MS (Fig S9). It has been demonstrated, previously, that acetyl-CoA binding happens on the cytosolic side and is optimal around pH 7.0-8.0, and the acetyl transfer activity happens on the luminal side and is optimal at pH 5.5-6.0. The K_m_ of *N*-acetyltransferase reaction in the presence of acetyl-CoA as the substrate is in the 0.2-0.6 mM range at pH 5-5.8, and 2.5-20 μM range at pH 7.0, suggesting that HGSNAT has a high affinity for acetyl-CoA at pH ≥7.0 (Bame & Rome, 1985, 1986a; Meikle *et al*, 1995). The transfer of the acetyl group from acetyl-CoA to the terminal non-reducing amino group of α-D-glucosamine is believed to be catalyzed by H269 located on the luminal entrance of ACOS (Fig 3A and Fig S5). It is unclear if H269 gets acetylated and forms a stable acetylated intermediate in the process of acetyl transfer. However, for the H269 to be acetylated or for it to transfer the acetyl group to its putative acceptor, protonation of the imidazole sidechain amine group is necessary. The catalytic H269 is primarily uncharged at pH 7.5, and the acetyl-CoA remains an acetate ion. The carboxamide group of N258 can act as both an acceptor and donor in a hydrogen bond. In our structure, ACOS is more accessible from the luminal side than the cytosolic entrance. We note that the acetyl-CoA is hydrogen-bonded to N258 (Fig 3A and Fig S5D). Based on these observations, we conclude that the structure we have is of an acetyl-CoA primed HGSNAT waiting for the protonation of H269 and the availability of an acetyl group acceptor to carry out acetyltransferase reaction (Fig 5A). Because the cytosolic pH is favorable for acetyl-CoA binding and the protein we purified endogenously pulled down acetyl-CoA with it, we believe this is the most stable conformation of HGSNAT in the absence of an acetyl group acceptor. Furthermore, in our FSEC-based analysis, we noticed that H269A substitution did neither alter the expression nor stability of HGSNAT (Figs S8A, S8D, and S8J). However, the N258I mutation completely destabilized HGSNAT (Figs S8A and S8C). We believe N258I instability to be a consequence of lack of substrate binding at the catalytic core, as we see a greater role for N258 in acetyl-CoA binding as opposed to H269 as per our structure (Figs 3A and S5).

**Figure 5:**
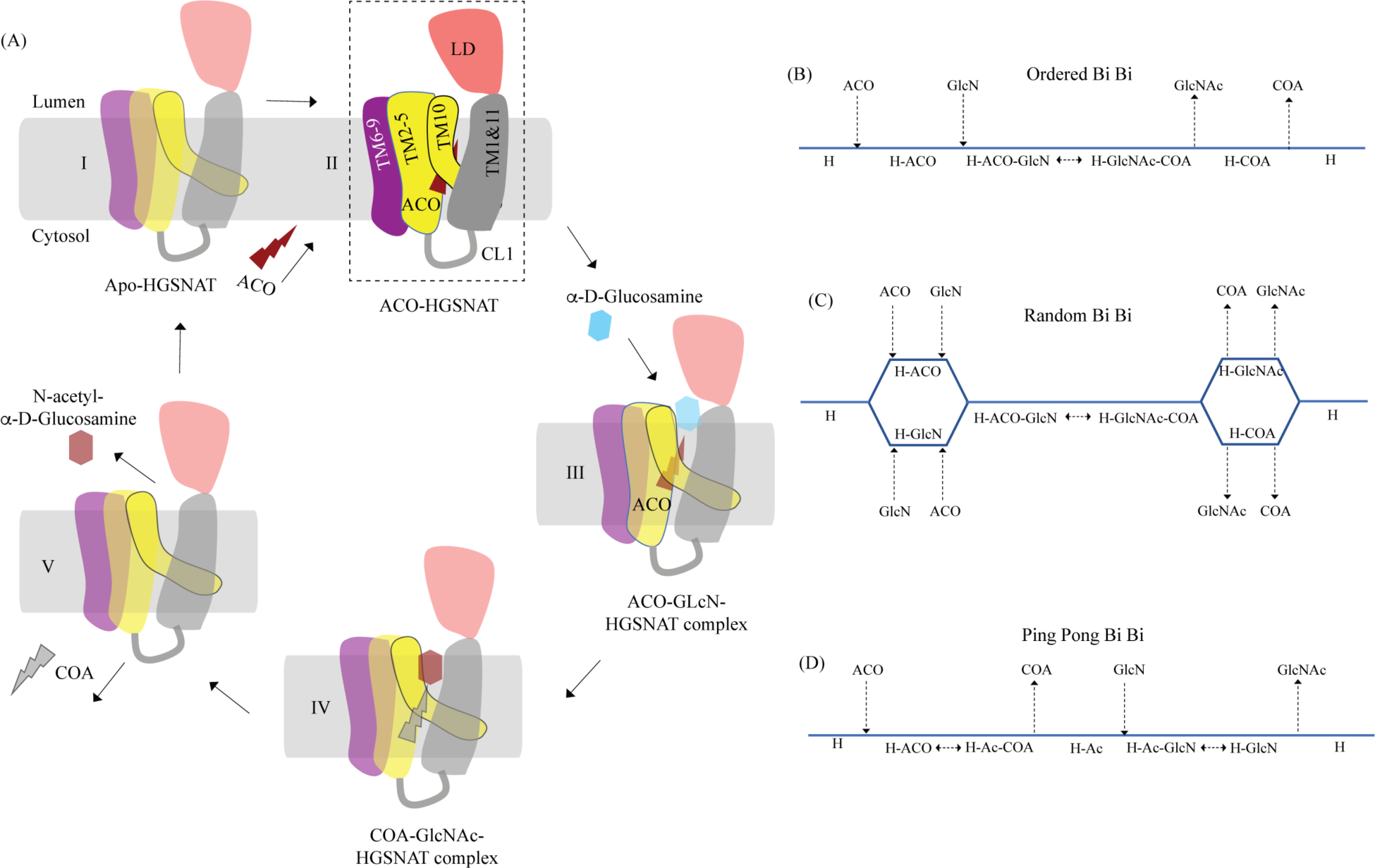
Proposed mechanism of acetyl transfer by HGSNAT. **(A)** HGSNAT (I) catalyzes a bisubstrate reaction of transferring acetyl group from cytosolic acetyl-CoA (ACO, red lightning) to terminal non-reducing α-D-Glucosamine (GlcN, blue hexagon) of luminal heparan sulfate (III and IV). After the acetyl group transfer, COA (gray lightning) and acetylated glucosamine (GlcNAc, red hexagon) are believed to be released to cytosol and lumen respectively (V). Depending on the order of binding and release of substrates and products, enzyme-catalyzed bisubstrate reactions could either be sequential reactions **(B** and **C)** or ping pong reactions **(D)**. The mechanism of reaction catalyzed by HGSNAT has been a longstanding debate. We believe that the acetyl-CoA bound HGSNAT structure presented in this work (II, dashed box) is in a cofactor primed conformation which could proceed by any of the bisubstrate reaction mechanisms shown in **B-D**. The function of LD is unclear, and we believe it plays essential role in recognition of substrate and its positioning at the active site.

## Discussion

Direct involvement of acetyl-CoA in the heparan sulfate degradation was shown in the early 1980s, by incubating the extracted intact lysosomes with radiolabeled acetyl-CoA and by subsequently monitoring the incorporation of radiolabeled acetate into lysosomal HS (Rome & Crain, 1981; Rome *et al*, 1983). Using purified lysosomal membranes, Rome and colleagues showed that the acetyl-CoA dependent *N-*acetyltransferase activity, which is required for acetylation of HS, resides on the lysosomal membrane. In these studies, they also suggested that the HS acetylation reaction occurs in two steps - acetylation of the lysosomal membranes and transfer of acetyl from the membranes to HS. By performing acetylation and acetyltransferase reactions at different pH, and in the presence of different amino acid modification reagents, they showed that the acetylation process is optimal at pH 7, acetyltransferase activity is optimal at pH 5.5, and the amino acid on the lysosomal membrane that gets acetylated is a histidine. They proposed that the *N*-acetyltransferase activity follows a Ping Pong Bi Bi mechanism of reaction, where acetyl-CoA from the cytosol acetylates the active site histidine of the *N*-acetyltransferase located on the lysosomal membrane, inducing a conformational change in the transmembrane enzyme that enables the access of active site histidine from the luminal side (Fig 5D). Due to the change in pH of the active site cavity, the acetyl-histidine interaction is destabilized, and the acetyl group is transferred to the α-D-glucosaminide on the luminal side (Bame & Rome, 1985, 1986a, b). Pshezhetsky and colleagues showed that acetylation of lysosomal membranes expressing *N*-acetyltransferase activity occurred even in the absence of the acetyl group acceptor, suggesting the formation of an acetylated enzyme intermediate (Ausseil *et al*, 2006; Durand *et al,* 2010). Here, we purified HGSNAT at a pH favorable for acetyl-CoA binding, but not acetyl transfer reaction. In our structure, we find that the HGSNAT ACOS is more accessible via the lumen as opposed to the cytosol and that the His269 of LL1 is not acetylated. Instead, we find an acetyl-CoA molecule in the ACOS, hydrogen bonded to N258 of TM2, a residue within a close vicinity (∼ 5 Å) of H269 (Fig 3 and Fig S5). We believe N258 holds onto acetyl-CoA and aids H269 in the acetyltransferase reaction, at low pH and in the availability of acetyl group acceptor.

Meikle and colleagues argued that the *N*-acetyltransferase reaction occurs via a random-order mechanism (Fig 5C). They demonstrated that both *N*-acetyl-α-D-glucosamine and COA are required for the reverse reaction, suggesting that the formation of a ternary complex of acetyl-CoA, α-D-glucosamine, and the enzyme enables the reaction to proceed in a single step without the requirement of an acetylated-enzyme intermediate. They also report two K_m_ values, that differ by 5-10 times, for both acetyl-CoA and glucosamine substrates, suggesting that both protomers, perhaps, bind to the substrates at varying affinity and only one of the protomers is catalyzing the acetyl transfer at any given point of time (Meikle *et al,* 1995). Mahuran and colleagues showed that acetylated enzyme intermediate could not be identified by affinity pulldown of purified transmembrane *N*-acetyltransferase after incubating it with [^3^H] acetyl-CoA and proposed that the formation of a stable acetylated enzyme intermediate is not required for the reaction to proceed (Fan *et al,* 2011). Our structure does not provide evidence of H269 acetylation (Fig 3). However, we observe that acetyl-CoA binds tightly to HGSNAT endogenously resulting in a stable HGSNAT-acetyl-CoA intermediate that stayed as an intact complex all through the affinity purification, size-exclusion chromatography, and cryo-EM sample preparation steps even without the addition of exogenous acetyl-CoA to any of the buffers (Fig S9). Perhaps, this tight endogenous binding of HGSNAT is the reason why Mahuran and colleagues could not observe labeling of purified transmembrane *N*-acetyltransferase by [^3^H] acetyl-CoA. The structure that we obtained does not by itself settle the debate of the mechanism of HGSNAT mediated acetylation. Our structure is in a conformation that could proceed through any of the three enzyme-catalyzed bisubtrate reaction mechanisms for acetyl transfer (Fig 5). While we were revising our manuscript, another group published structures of apo HGSNAT and HGSNAT-ACO, HGSNAT-COA-NAG complexes. All these structures were obtained at pH 8.0 in 1% GDN and revealed no drastic conformational isomerization in the protein except for localized motion in TM2 and TM3 and residues at ACOS (Xu *et al*, 2024). Xu and colleagues do not show if the order of binding of the substrate is crucial for catalysis, and nor do they disprove the existence of acetylated enzyme intermediate. We believe structures of HGSNAT at lysosomal luminal pH in the presence and absence of an acetyl group acceptor will demonstrate not only if the enzyme gets acetylated during the reaction but also show if there are pH driven conformational changes within the protein. It is also likely that the choice of detergent used determines the conformational isomerization. So, obtaining structures in membrane mimetics like nanodiscs in presence of native HGSNAT lipids could provide a clearer insight into HGSNAT catalyzed reaction mechanism.

HGSNAT is believed to be produced as a pro-protein that gets proteolytically processed into its mature form where two fragments of unequal sizes, the α-HGSNAT and β-HGSNAT, continue to stay together because of a disulfide bond mediated interaction between C123 (β6) in the N-terminal luminal domain (LD) of the α-HGSNAT and C434 in the LL3 of the *C*-terminal transmembrane domain (TMD) of β-HGSNAT (Durand *et al,* 2010). However, according to the structure, these amino acids are ∼ 34 Å apart making such a bond impossibility in the acetyl-CoA bound HGSNAT conformation reported here (Fig 1A, 1D and 2B). Moreover, we notice that C434 is disulfide bonded to C415 on LL3. C123 of β6 is predicted to form a disulfide bond with C151 of β8, and we do not see density to model such a disulfide. A bond between C123 of LD and C434 of LL3 could be a possibility only if there is a drastic conformational change during that brings LD closer to the LL3. Thus, we believe that the α-HGSNAT and β-HGSNAT stay together because of a series of hydrogen bonds, dipole-dipole and hydrophobic interactions between β2-β3 turn, β4-β5 turn, LL1, LL3, and LL4 (Fig 2 and Fig S6).

The protease that cleaves HGSNAT into α-HGSNAT and β-HGSNAT is unknown, and the site for proteolysis is unclear. Fan et al, proposed that the site of proteolysis is between amino acids N144 and G145 in the LD (Fan *et al,* 2011). N144-G145 is located on the β7-β8 turn of the LD, a region poorly resolved in our structure (Fig 1A and Fig 2A). Although proteolysis at this position will cleave the LD into two parts (β1-7 and β8), the LD should remain attached to TMD because of extensive network of interactions along the LD-TMD interface (Fig 2B and Fig S6). Durand et al, proposed that the site of proteolysis is between the end of the luminal domain and beginning of LL1 (Durand *et al,* 2010). Sequence-based protease site prediction in Procleave and Prosper servers, using the sequence between β8 of LD and LL1, suggests multiple potential sites of proteolysis, for example, L154-A155 at the C-term of β8, S162-N163 in the loop connecting LD to TM1, F170-L171 at the N-term of TM1, L187-S188 and W231-R232 at the N- and C-term of the CL1 (Li *et al*, 2020; Song *et al*, 2012). It is hard to speculate which of these are most probable, as the lysosomes are rich in aspartic, serine, and cysteine proteases, including intra-membrane proteases that are involved in protein degradation and proprotein processing. In addition, some matrix metalloproteinases have also been shown to act in intracellular compartments and have been implicated in MPS diseases (Batzios *et al*, 2012; Jobin *et al*, 2017; Muller *et al*, 2012; Schroder & Saftig, 2016; Turk *et al*, 2000). The recombinant HGSNAT that we produced is a non-proteolyzed form of HGSNAT, as we see a band corresponding to full-length HGSNAT in our SDS-PAGE analysis and a peak corresponding to an intact full-length dimer in our FSEC analysis. HGSNAT is encoded by 18 exons, where the first 6 exons encode an N-terminal luminal domain (LD) & TM1, which are only present in the metazoans. The remaining 10 TMs and the C-terminus are well conserved in archaea, bacteria, and plants (Fan *et al,* 2006; Hrebicek *et al,* 2006). Based on the evolutionary sequence and structure conservation and conformational flexibility observed in CL1, we speculate that LD & TM1 together form α-HGSNAT and TMs 2-11 form β-HGSNAT (Fig 2A, Fig 4A and Fig S4A).

There are no known homologs of the LD, and even the strand arrangement in two sheets appears to be rare (Koch *et al*, 1992). A structure-based search in the Dali server lists hits with <15% identity, with immunoglobulin-like & transthyretin folds as top hits (Table S1) (Beale *et al*, 2015; Felisberto-Rodrigues *et al*, 2011; Parker *et al*, 2021; Rajasekar *et al*, 2019). Unlike the LD, a typical immunoglobulin-like fold is a disulfide-containing β-sandwich made of 7-9 β-strands arranged in two anti-parallel β-sheets, with a conserved core formed by four strands β2, β3, β5, & β6 (Fig S4C and S4D). In the Ig-like fold, the β2 and β5 strands are in one sheet, and the β3 and β6 strands are in the second one, with β2 and β6 being linked by a disulfide bond. The remaining 3-5 strands comprise the remaining variable region of the Ig-like fold (Bork *et al*, 1994; Chidyausiku *et al*, 2022). Although the % identity of LD and the transthyretin fold containing hits is <15%, the strand composition of the two sheets that make the β-sandwich is identical. However, a typical transthyretin fold contains a conserved short α-helix between β5 & β6 and often does not contain disulfide bonds (Fig S4C, D) (Hornberg *et al*, 2000). Another β-sandwich that shows a similar secondary structure assignment as LD is the type-II C2 domain, where the top sheet is made of β1, β4, β7, & β8 strands while the bottom sheet is made of β2, β3, β5, & β6 (Hirano *et al*, 2019; Kretsinger *et al*, 2013). However, both sheets in the type-II C2 domain are anti-parallel, and the arrangement of the top strand is β4-β1-β8-β7. In LD, one sheet is anti-parallel, and the other is a mixed sheet where the arrangement in the top sheet is β4-β1-β7-β8 (Fig S4C and S4D). Because of these non-trivial dissimilarities of LD with three major types of β-sandwiches, we believe that LD is a new fold of β-sandwich that could be best described as a ‘transthyretin-like’ fold. In fact, two proteins in the top hits from a structure-based similarity search on Dali server show the same sheet arrangement as LD (Table S1 and Fig S4D). One of these proteins is AlgF (PDB ID: 6CZT), an adaptor protein from Pseudomonas believed to be involved in O-acetylation of alginate exopolysaccharides and the other is an SPH (self-incompatibility protein homolog) domain (PDB ID: 6G7G) that is described as transthyretin-like and is believed to be a plant secreted protein involved in cell death (Fig S4D).

The function of LD remains unknown. DeepSite predicts the presence of two ligand binding sites in HGSNAT, one that overlaps with the ACOS and the other in the cavity between the two sheets of LD (Fig S5B) (Jimenez *et al*, 2017). However, it is unclear if LD binds to a ligand and/or a metal ion. It has been shown that the *N*-acetyltransferase activity of HGSNAT is enhanced in the presence of anionic phospholipids (Fan *et al,* 2011). However, we expect lipids to bind towards the cytosolic side at the dimer interface. We see unexplained density in these regions that could be best explained as ordered lipids or digitonin (Fig S7). A recent report of a computational modeling and molecular dynamics simulation study conducted on a model of OafB, a bacterial O-antigen modifying transmembrane acetyltransferase of the ATAT family within the TmAT superfamily, showed that the periplasmic SGNH domain undergoes large conformational changes and aid in O-antigen acetylation (Newman *et al,* 2023). OafB has two domains – the transmembrane acetyltransferase-3 domain (AT-3) and periplasmic SGNH domain. Although the domain organization is similar to HGSNAT, there are marked differences in the structure (Fig S4B). AT-3 domain and HGSNAT TMD are not homologous, even though they share a similar architecture of the ACOS. The LD of HGSNAT is different from the SGNH domain, in the sense that LD is a β-sandwich at the N-terminus of the protein while SGNH is a α-β-α sandwich with a single β-sheet sandwiched between two helical domains at the C-terminus of the protein. There are non-trivial differences between the predicted structures of ATAT and YeiB family members. However, it is possible that the TMDs and the extra-membranous domains function similarly. It is likely that LD of HGSNAT binds to HS, stabilizing the terminal non-reducing sugar of HS near the luminal side of ACOS for catalysis and remains to be structurally investigated. In fact, it has been shown that as the size of acetyl group acceptor was increased in an acetyltransferase reaction from mono-to di-to tetra-saccharide, the K_m_ values decreased from 0.6 mM to 7 μM (Meikle *et al,* 1995).

The high-resolution structure of HGSNAT reported in this study heralds a new beginning for structural exploration of the TmAT superfamily. The conformation of HGSNAT observed in our cryo-EM studies does not put to rest the debate on the kind of bi-substrate reaction mechanism that HGSNAT follows to catalyze the acetyl transfer. However, our structure underscores the requirement to characterize the remaining states of the enzymatic reaction. The molecular basis we delineated for the mutation-induced dysfunction in HGSNAT will serve as a blueprint for the structure-based design of novel therapeutic modulators that could rescue the function of milder variants.

## Materials and Methods

### Cloning and site-directed mutagenesis

The codon-optimized gene encoding the isoform 2 of full-length human HGSNAT was synthesized by GenScript. The synthesized gene was then cloned into the pEG BacMam expression vector (Addgene plasmid # 160683) between EcoRI and NotI restriction sites, to be expressed via baculoviral transduction in HEK293S GnTI^-^ cells (ATCC # CRL-3022) as a fusion protein containing an N-terminal Strep-tag-II-GFP. The integrity of the clone was confirmed by Sanger sequencing (Plasmidsaurus, OR). HGSNAT variants discussed in the manuscript were prepared in the pEG BacMam expression vector background by GenScript.

### Cell culturing, transient transfection, and transduction

Adherent HEK293S GnTI^-^ cells were grown in Dulbecco’s Modified Eagle Medium (DMEM, Gibco) supplemented with 10% fetal bovine serum (FBS, Gibco) at 37°C. Immediately preceding transfection, the cells were washed with 1X PBS (Gibco) and supplied with fresh pre-warmed DMEM containing 10% FBS. 1x10^6^ cells were transfected with 1μg DNA using TurboFect (Thermo Fisher Scientific) suspended in serum-free DMEM as suggested by the manufacturer’s protocol. The transfected cells were grown at 37°C and 5% CO_2_ for 8-10 hr. The cell-culture media was then replaced with fresh pre-warmed DMEM containing 10% FBS and 10 mM of sodium butyrate, and the cells were grown at 32°C and 5% CO_2_ for an additional 24-36 hr, before harvesting. All transfections were done at 80% confluency. Transfected cells were used to screen for ideal expression and purification conditions.

Baculovirus preparation was done as described by Goehring and colleagues (Goehring *et al,* 2014). Briefly, DH10Bac cells (Thermo Fisher Scientific) were transformed with an HGSNAT expression vector, and lacZ^-^ colonies were selected on gentamycin-kanamycin-tetracycline LB agar plates for bacmid DNA isolation. 1x10^6^ adherent Sf9 cells (ATCC # 12659017) grown in serum-free Sf-900 III media (Gibco) were transfected with 1 μg of bacmid using Cellfectin II reagent (Gibco) per the manufacturer’s protocol. Transfected cells were grown at 27°C for 96 hr. The supernatant media from the cells was harvested, filtered through a 0.2 μm filter, and stored as P1 virus. 100 μL of P1 virus was added to 1L of Sf9 cells at a cell density of 1x10^6^/mL in serum-free Sf-900 III media. The cells were grown at 96 hr at 27°C while shaking at 120 rpm. The cells were spun down at 4000 xg for 20 min, and the supernatant media was filtered through a 0.2 μm filter and stored as P2 baculovirus. P2 baculovirus was used for large-scale transduction of HEK293S GnTI^-^ cells. Large-scale expression of HGSNAT was done by baculoviral transduction of HEK293S GnTI^-^ cells suspended in FreeStyle 293 expression media containing 2% FBS. Mammalian cell culture at a density of 3x10^6^ cells/mL was transduced using P2 baculovirus at a multiplicity of infection of 1.5-2 and was incubated on an orbital shaker at 37°C and 5% CO_2_ for 8-10 hr. The cells were supplemented with 10 mM sodium butyrate and were then incubated in a shaker at 32°C and 5% CO_2_ for an additional 38-40 hr. The cells were harvested by centrifugation at 4000 xg for 10 min, and the cell pellet was stored at -80°C until further use.

### Protein expression and thermostability analysis

To identify suitable conditions for large-scale expression and solubilization of HGSNAT, we employed fluorescence-detection size-exclusion chromatography (FSEC) (Kawate & Gouaux, 2006). Briefly, 100,000 transfected cells were solubilized in 500 μl of 1 % detergent, 25 mM Tris-HCl, pH 7.5, 200 mM NaCl, 1 mM PMSF, 0.8 μM aprotinin, 2 μg/mL leupeptin, and 2 μM pepstatin A at 4°C for 1 hr on an end-end rotator. After solubilization, the lysate was centrifuged at 185,000 xg for 1 hr at 4°C. The supernatant was filtered through 0.45 μm filter, and 100 μl filtrate was analyzed on a Superose 6 Increase 10/300 GL column (Cytiva Life Sciences) pre-equilibrated with 0.15 mM LMNG, 25 mM Tris-HCl, pH 7.5, and 200 mM NaCl. GFP fluorescence in the eluate was monitored by a fluorometer (Shimadzu scientific instruments) set at Ex/Em of 485/510 nm. To test the stability of HGSNAT in various conditions, the solubilized samples were heated at 55°C for 15 min, centrifuged at 10,000 rpm for 10 min, filtered through 0.45 μm filter, and were analyzed again on a Superose 6 Increase 10/300 GL column. To measure overall relative expression of mutants, 100,000 cells expressing the mutants were solubilized in 100 ul of 1% digitonin, 25 mM Tris-HCl, pH 7.5, 200 mM NaCl, 1 mM PMSF, 0.8 μM aprotinin, 2 μg/mL leupeptin, and 2 μM pepstatin A at 4°C for 1 hr on an end-end rotator. The solubilized lysates were transferred to Costar 96-well flat bottom clear plates without centrifugation, and GFP fluorescence was monitored at Ex/Em of 480/520 nm in SpectroMax M5 (Molecular Devices) microplate reader. For FSEC analysis, mutants were solubilized in 1% digitonin, but were also centrifuged and filtered before being analyzed on Superose 6 Increase 10/300 GL column. Thermal stability of the mutants was tested by heating solubilized lysates of the mutants and WT HGSNAT at 65°C for 15 min and analyzing them by FSEC.

### Purification of HGSNAT

Cell debris from the sonicated lysate of HEK293S GnTI^-^ cells expressing N-terminal GFP fusion of HGSNAT was removed by centrifugation at 2400 xg for 10 min. The supernatant from the low-speed centrifugation step was subjected to ultra-centrifugation at 185,000 xg for 1 hr at 4°C to harvest membranes. Membranes were resuspended using a dounce homogenizer in suspension buffer (25 mM Tris-HCl, pH 7.5, 200 mM NaCl, 1 mM PMSF, 0.8 μM aprotinin, 2 μg/mL leupeptin, and 2 μM pepstatin A). To this membrane suspension, an equal volume of 2% digitonin solution prepared in membrane suspension buffer was added. The membrane suspension was solubilized for 90 min. at 4°C. The solubilized membrane suspension was centrifuged at 185,000 xg for 1 hr at 4°C. The solubilized supernatant was passed through Strep-Tactin affinity resin (IBA Life Sciences) column at a flow rate of ∼0.3-0.5 ml/min. Affinity resin saturated with HGSNAT was washed with 5-6 column volumes of 0.5% digitonin, 25 mM Tris-HCl, pH 7.5, and 200 mM NaCl. The bound protein was eluted using 5 mM D-desthiobiotin prepared in the wash buffer. The purity and homogeneity of purified HGSNAT was confirmed by performing FSEC and SDS-PAGE. The amino acid sequence of purified HGSNAT was verified by peptide mass fingerprinting of the bands on SDS-PAGE. For FSEC experiments, ∼10 μL of elution fractions were loaded onto a Superose 6 Increase 10/300 GL column pre-equilibrated with 0.15 mM LMNG, 25 mM Tris-HCl, pH 7.5, and 200 mM NaCl. The elution fractions containing homogeneous and pure protein were pooled and concentrated to 2 mg/mL. The concentrated protein was further purified by size-exclusion chromatography (SEC) using Superose 6 Increase 10/300 GL column pre-equilibrated with 0.5% digitonin, 25 mM Tris-HCl, pH 7.5, and 200 mM NaCl. HGSNAT corresponding to the dimeric peak from the SEC step was pooled and concentrated to 0.9 mg/mL for cryo-EM sample preparation.

### LC-MS analysis of purified HGSNAT

To identify the endogenously bound acetyl-CoA in recombinant HGSNAT, the purified protein was loaded onto a Chromolith RP-18 endcapped column (4.6 mm X 100 mm; Supleco) operated at 1 ml/min flow rate and 40°C. Acetyl-CoA was eluted by a gradient of mobile phase A (5mM ammonium formate) and mobile phase B (10-90% acetonitrile). The eluted samples were analyzed by a Thermo TSQ-Quantis (Thermo Scientific) in positive mode at 5.5kV capillary voltage while scanning at 1000 Da/sec from 200-1000 m/z in single reaction monitoring (SRM) mode. The data was analyzed using Thermo FreeStyle 1.6 (Thermo Scientific), using a SRM filtering for acetyl-CoA precursor (810.1 m/z) and product (303.1 m/z). To monitor the specific endogenous binding of acetyl-CoA to HGSNAT, we also purified apo HGSNAT from the membrane suspension dialyzed in membrane suspension buffer for 4 days at 4°C. Apo HGSNAT was processed for LC-MS analysis in the same fashion as HGSNAT-ACO complex.

### Cryo-EM sample preparation and data collection

UltrAuFoil holey-gold 300 mesh 1.2/1.3 μm size/hole space grids (UltrAuFoil, Quantifoil) were glow discharged, using a PELCO glow discharger, for 1 min at 15 mA current and 0.26 mBar air pressure and were immediately used for sample vitrification. 2.5 μL of HGSNAT at 0.9 mg/ml was applied to the glow-discharged grids, and subsequently blotted for 2s at 18°C and 100% humidity using a Vitrobot (mark IV, ThermoFisher Scientific). Without any wait time, the grids were plunge-frozen in liquid ethane. Movies were recorded on a Gatan K3 direct electron detector in super-resolution counting mode with a binned pixel size of 0.85 Å per pixel using Serial EM on a FEI Titan Krios G4i transmission electron microscope operating at an electron beam energy of 300 KeV with a Gatan Image Filter slit width set to 20 eV. Exposures of ∼2s were dose fractionated into 50 frames, attaining a total dose of ∼50 *e*^-^ Å^-2^, and the defocus values were varied from -1.0 to -2.5 μm.

### Cryo-EM data processing

All image processing was done using CryoSPARC (v 4.2.1) unless specified otherwise (Punjani *et al*, 2017). Briefly, motion correction was carried out on the raw movies, and subsequently, the contrast transfer function (CTF) was estimated using patch-CTF correction on dose-weighted motion-corrected averages within cryoSPARC (Fig S2). A total of 15,067 micrographs were collected, of which 10,394 were selected for further processing based on the CTF fit. Both blob picking and template picking were used to pick particles, and duplicates were removed. Particles were extracted initially using a box size of 288 pixels. A subset of the particles picked by the blob picker was used to generate a low-resolution *ab initio* reconstruction, which was later used as a reference for iterative 3D classification and heterogenous refinement. The pooled particles were subjected to multiple rounds of reference-free 2D classification and heterogenous refinement to get a cleaned subset of particles which resulted in classes with recognizable features. The cleaned particles were re-extracted with a box size of 360 pixels, and a stack of 94,875 good particles was selected and subjected to an initial round of NU refinement using an *ab initio* model as a reference (Punjani *et al*, 2020). 86,500 particles resulted in a 3.89 Å map, which had well-resolved luminal domain architecture, and visible secondary structure features of TM helices. To improve the resolution further, only the particles belonging to micrographs with CTF fit <4 Å were selected for subsequent refinements. Both C1 and C2 symmetry-imposed maps overlapped well, and we used C2 symmetry in the final rounds of processing (Fig S2 and S3C). A final subset of 57,739 particles was used for NU-refinement, yielding a 3.26 Å map (Fourier shell coefficient (FSC) = 0.143 criterion). Symmetry expansion and local refinement with a focused mask on LD and TMD domain was performed to improve local density in these domains. The map obtained by symmetry expansion was only used to assess the fit of the model to density. A composite map generated by combining these local maps was used to finalize the fit of the side chains during model building and refinement (Fig S3D). Local resolutions were estimated using the RESMAP software (Kucukelbir *et al*, 2014).

### Model building, refinement, and structure analysis

ModelAngelo was used to build HGSNAT isoform 2 into the final map, and final refinement was performed in Phenix (Jamali *et al,* 2023; Liebschner *et al*, 2019). Coot was used to rebuild certain portions of the protein and to analyze the Ramachandran outliers (Emsley *et al*, 2010). ChimeraX was used for making the figures (Meng *et al*, 2023). Ligand binding site prediction on HGSNAT was performed using MOLEonline and DeepSite (Jimenez *et al,* 2017; Pravda *et al,* 2018). To analyze the effect of missense mutations on HGSNAT and to calculate free energies indicating the relative mutation-induced destabilization, we used FoldX (Schymkowitz *et al,* 2005).

## Acknowledgements

We thank the University of Michigan (UM) cryo-EM facility staff for assistance with cryo-EM data collection, and Drs. Michael Cianfrocco, Farzad Jalali-Yazdi, Jonathan Coleman, and Prashant Rao for helpful comments on cryo-EM analysis. UM cryo-EM facility receives generous support from UM BSI and UM LSI. We thank all the members of Mosalaganti lab and Baldridge lab for their educative discussions on our work. This work was supported by the start-up funding provided to SM by the University of Michigan. SM is a recipient of the NIH Director’s New Innovator Award (DP2) (1DP2GM150019-01).

## Author contributions

VN designed the project, prepared the sample, collected the cryo-EM data, and wrote the first draft of the manuscript. AK, VN, and SM processed and analyzed the cryo-EM data. VN and AK built the model and made the figures. AK deposited the map and coordinates. JKR performed the experiments with HGSNAT variants and prepared samples for LC-MS. VN, AK, JKR, and SM contributed to the editing and preparation of the final draft of the manuscript.

## Conflict of interest

The authors declare no conflict of interest.

## Data availability

The data presented in this manuscript are available upon request. All correspondence for materials and data presented in this paper should be addressed to. The cryo-EM maps and 3D coordinates of HGSNAT have been deposited in the Electron Microscopy Data Bank (EMDB) and Protein Data Bank (PDB) under the accession codes EMD-41620 and PDB-8TU9. All the half-maps and the masks used for refinement have also been deposited in the EMDB under the same accession code.

## Supplementary file

### Supplementary figures

**Figure S1:**
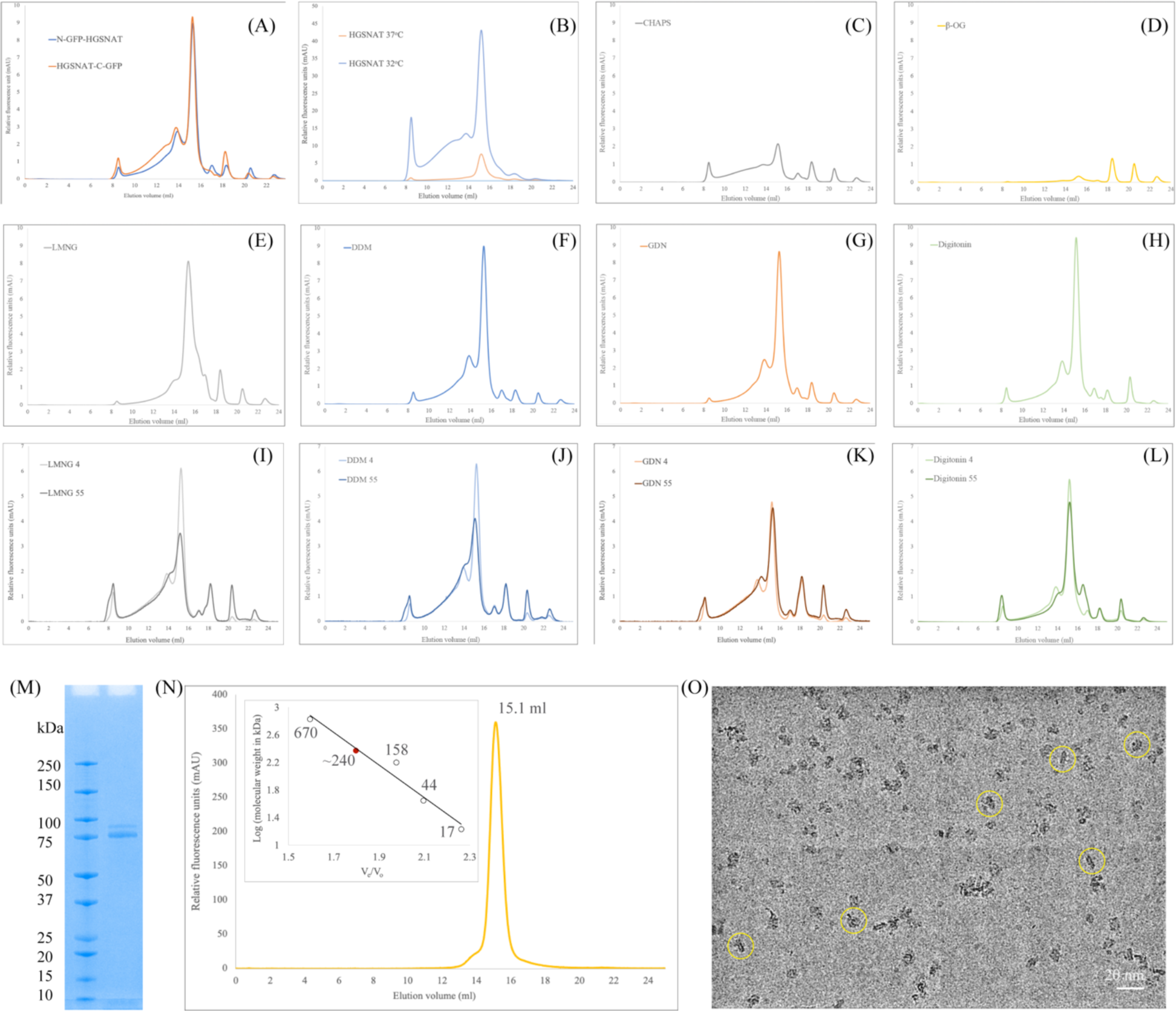
Purification of HGSNAT. **(A)** Comparison of expression of N- and C-terminal GFP fusions of HGSNAT in HEK293S GnTI^-^ cell lysates, solubilized in 1% DDM. **(B)** Comparison of relative overexpression of N-GFP-HGSNAT in cultures grown at 37°C and 32°C, post-transduction. **(C-H)** Relative solubility and homogeneity comparison in 1% of CHAPS, β-OG, LMNG, DDM, GDN, and digitonin, respectively, prepared in 25 mM Tris-HCl, pH 7.5, 200 mM NaCl, 1 mM PMSF, 0.8 μM aprotinin, 2 μg/mL leupeptin, and 2 μM pepstatin A. **(I-L)** Comparison of relative thermal stability of detergent solubilized HGSNAT in 1% of LMNG, DDM, GDN, and digitonin respectively. Samples analyzed after heat treatment at 55°C for 15 min have been marked with a suffix 55, and samples stored in cold room are marked with a suffix 4. **(M)** SDS-PAGE (12%) showing purity and monomeric molecular weight of HGSNAT. Although, monomeric molecular weight is ∼ 100 kDa, the full-length GFP fusion of HGSNAT, like most eukaryotic membrane proteins, displays anomalous electrophoretic mobility and runs around 75 kDa. **(N)** Intrinsic tryptophan fluorescence size-exclusion chromatogram of purified HGSNAT analyzed on Superose 6 Increase 10/300 GL column at 0.5 ml/min flowrate in LMNG-based FSEC running buffer. Red dot on the standard plot (log of protein molecular weight (kDa) vs. ratio of the elution volume to the void volume (V_e_/V_o_) indicates that recombinant N-GFP-HGSNAT elutes at 15.1 ml corresponding to a dimer of ∼ 240 kDa. **(O)** Representative micrograph imaged on Titan Krios using UltrAuFoil holey-gold 300 mesh 1.2/1.3 μm grid of vitrified N-GFP-HGSNAT at 0.9 mg/ml. The protein (yellow circles) distribution on grids, along with SDS-PAGE and size-exclusion chromatogram shows a monodisperse sample preparation.

**Figure S2:**
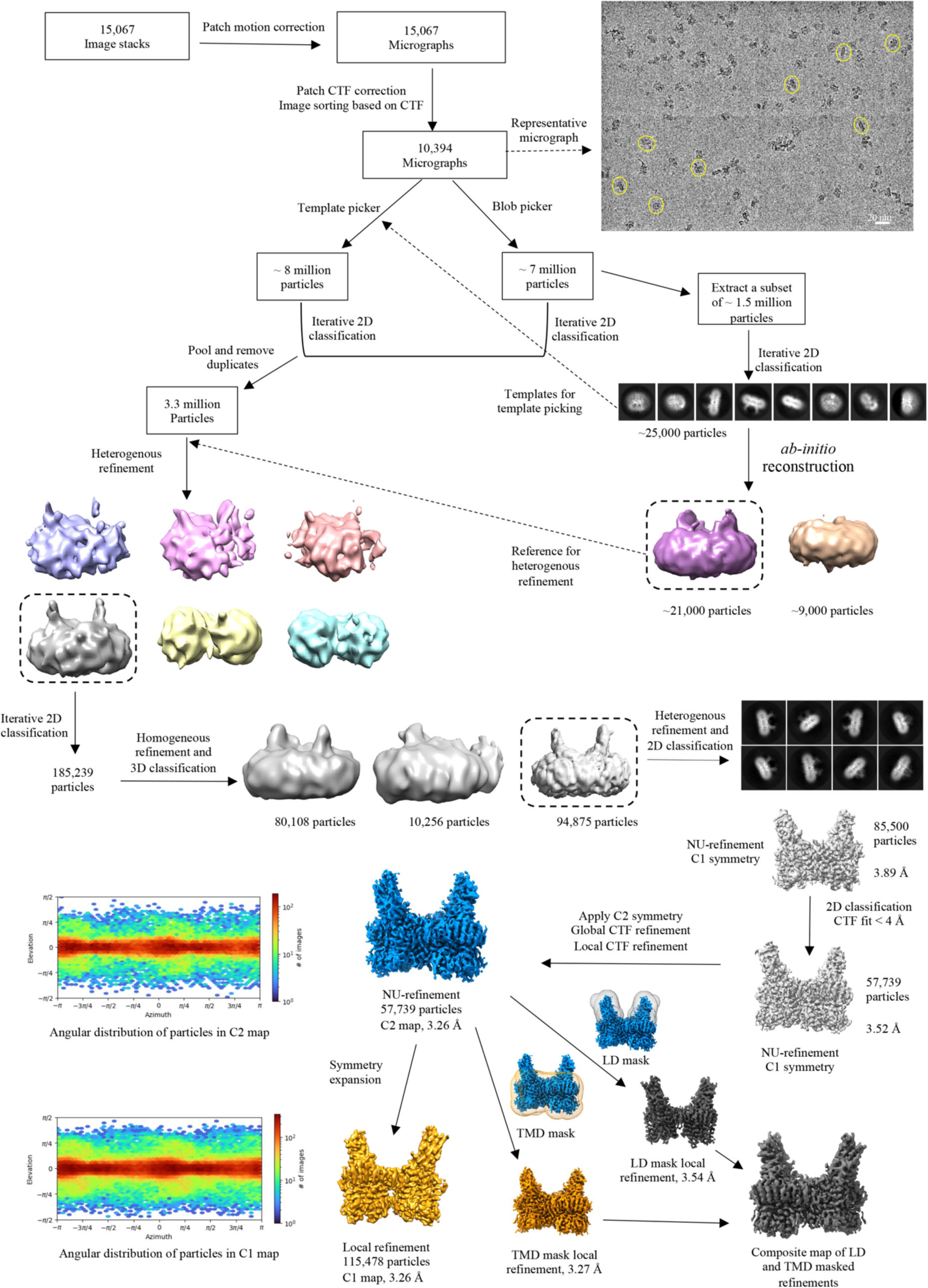
Cryo-EM data processing workflow. The data was entirely processed in cryoSPARC. A representative motion-corrected micrograph with single HGSNAT particles (yellow circles) is highlighted. A subset of (1.5 million) particles picked by blob picker were extracted and cleaned by 2D classification to generate 2D templates for template-based picking and an ab-initio volume to be used as reference input for subsequent heterogenous refinement jobs (dashed arrows). Particles picked using template picker and blob picker were individually cleaned by 2D classification to remove obvious junk particles and then were pooled and duplicates were removed for sorting by heterogenous refinement, and iterative 2D and 3D classification. Throughout the processing workflow classes with most well-resolved luminal domain was used as input references for the subsequent steps of processing (highlighted by a dashed boxes). A resultant stack of 85500 particles (representative 2D classes highlighted) was further cleaned up based on CTF fit (<4 Å) to end up with a final particle stack of about 57000 particles. C2 symmetry was applied at this stage and non-uniform and CTF refinements were performed to yield a C2 map at 3.26 Å. This map was used for model building and analyzing the structure of HGSNAT. A C1 map was generated by symmetry expansion of the final particle stack followed by local refinement to compare the quality of data with and without C2 symmetry application. Local refinements with masks focused on LD and TMD domain were performed separately, and a composite map was generated by combining these local refined maps to improve the density in these regions. The composite map was only used to finalize the fit of LBD side chains in the final model.

**Figure S3:**
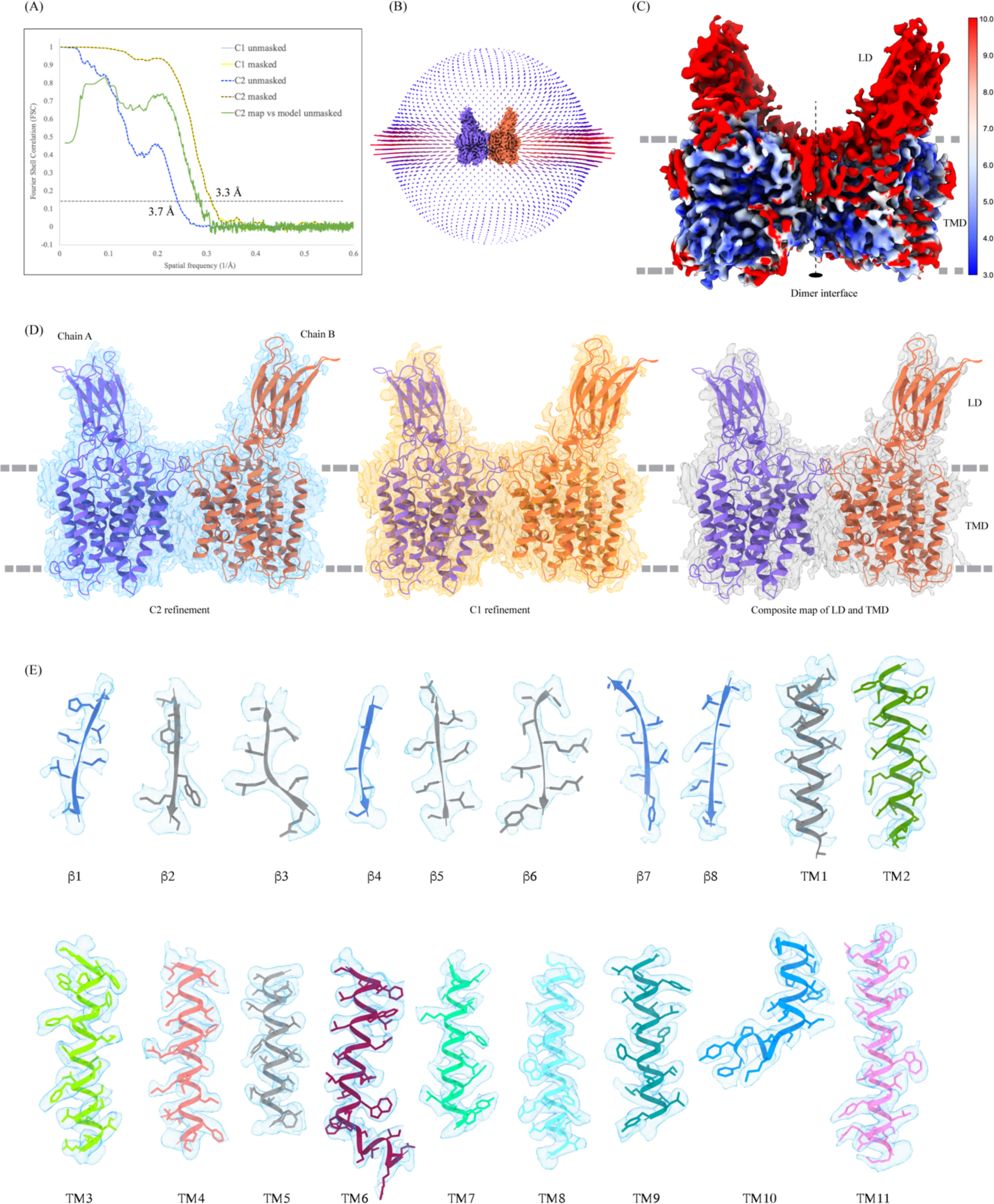
Cryo-EM data quality, reconstruction, and model building. **(A)** FSC curves for cross-validation. The final masked HSGNAT (C1: light yellow; C2: dark yellow dashed) and unmasked (C1: light blue; C2: dark blue dashed) refinement maps. Model vs. final C2 map unmasked (green). Gray and black dashed lines indicate FSC=0.143 and FSC=0.5 thresholds respectively. FSC curves were calculated using Mtriage in Phenix. **(B)** Angular distribution of particles used in the final reconstruction. **(C)** C2 map colored by estimated local resolution. **(D)** HGSNAT modelled by ModelAngelo into the C2 map (blue). The fit of the same model in C1 (orange) and composite map (gray) of LD and TMD created in ChimeraX. All maps are displayed at level 0.21 in ChimeraX. **(E)** Cryo-EM density of all the secondary structure elements, β1-β8 and TMs 1-11, shown in light blue (display level between 0.18-0.25 in ChimeraX). Side chains for almost all the elements could be modeled unambiguously into the density. At places with missing density, the side chains were trimmed to Cβ.

**Figure S4:**
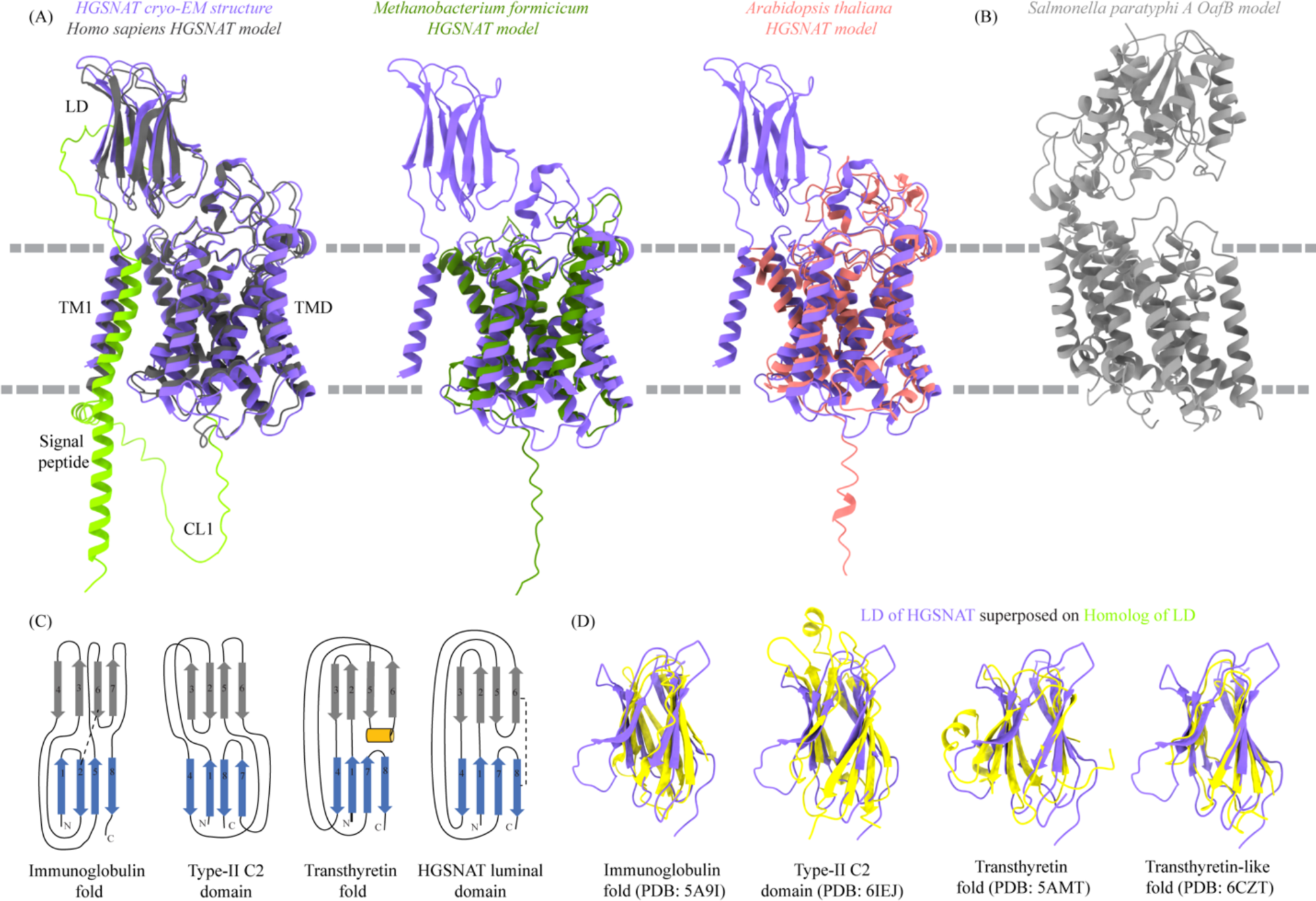
Homologs of HGSNAT. **(A)** Superposition of HGSNAT cryo-EM structure (purple) with the AlphaFold models of human (Uniprot: Q68CP4, dark gray), *Methanobacterium formicicum* (Uniprot: K2QAW2, green), and *Arabidopsis thaliana* (Uniprot: A0A5S9Y8V3, pink) HGSNATs. Cα RMSDs of the superpositions are 1.34 Å, 1.17 Å, and 1.13 Å respectively, suggesting a conserved HGSNAT fold across different kingdoms. AlphaFold model of HGSNAT shown here is of isoform 1, that has extra 28 residues on the N-term as compared to isoform 2. The structure is of isoform 2. The cryo-EM density did not allow modeling of residues upstream of β1 on the N-terminus and CL1, which have been highlighted yellow in the AlphaFold model. **(B)** AlphaFold model of acetyltransferase model of *Salmonella paratyphi* A OafB, an O-antigen modifying transmembrane acetyltransferase of the ATAT family within the TmAT superfamily (Uniprot: A0A0H2WM30). Despite predicted to be in the same superfamily as HGSNAT, a meaningful alignment and similarity to HGSNAT was not observed, highlighting the diversity of membrane bound acetyltransferases. **(C)** Comparison of topologies of immunoglobulin (Ig) fold, type-II C2 domain, and transthyretin fold with LD of HGSNAT. Strands in two sheets are colored blue and gray and conserved helical turn in transthyretin fold is shown in orange. Conserved disulfides are shown as dashed lines. **(D)** Superposition of structures of Ig fold (PDB: 5A9I), type-II C2 domain (PDB: 6IEJ), transthyretin fold (PDB: 5AMT), and transthyretin-like domain (PDB: 6CZT) onto LD of HGSNAT. Based on the topology and structure superposition, it appears that LD is transthyretin-like domain.

**Figure S5:**
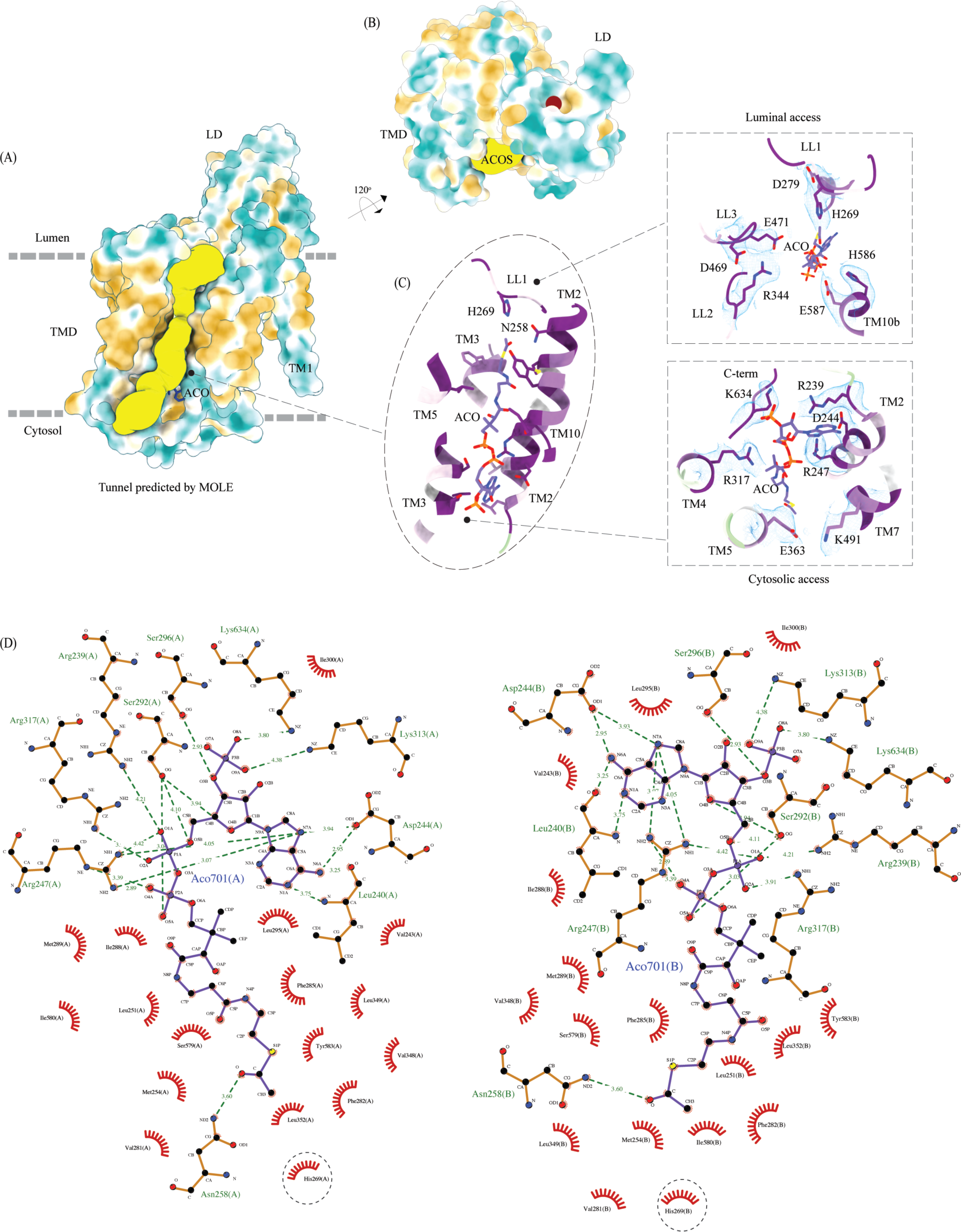
Ligand binding sites of HGSNAT. **(A)** Surface representation of HGSNAT (chain A), with hydrophobic and hydrophilic amino acids colored in orange and cyan respectively. Predicted acetyl-CoA access tunnel (yellow) by MOLEonline with a probe radius of ∼ 1.5 Å (Pravda *et al,* 2018). ACO bound at HGSNAT is shown in blue. It is apparent that the nucleoside head group and the acetyl group interact with hydrophilic residues and the pantothenate group is supported by hydrophobic residues. **(B)** Ligand binding site on LD (maroon sphere) predicted by DeepSite (Jimenez *et al,* 2017) **(C)** ACOS color coded based on the evolutionary sequence conservation scores obtained from ConSurf server. In the insets are the integral salt-bridges of the luminal (top) and cytosolic (bottom) access of ACOS. The cryo-EM density for the salt-bridges is shown in blue (display level 0.22 of the C2 refine map in ChimeraX). **(D)** 2D depiction of the network of interactions of ACO modeled at chain A (left) and chain B (right) with HGSNAT residues that lie <4.5 Å away from ACO, generated in LigPlot+. Hydrogen bonds are depicted by dashed lines, and residues that are involved in hydrogen bonds with ACO are shown as ball & stick models. Non-bonded contacts are indicated as eye lashes. The predicted active site H269 is highlighted by dashed circle. In our structure N258 forms weak hydrogen bonds with the acetyl group of ACO. We believe that N258 holds onto ACO until H269 is protonated and ready for catalysis.

**Figure S6:**
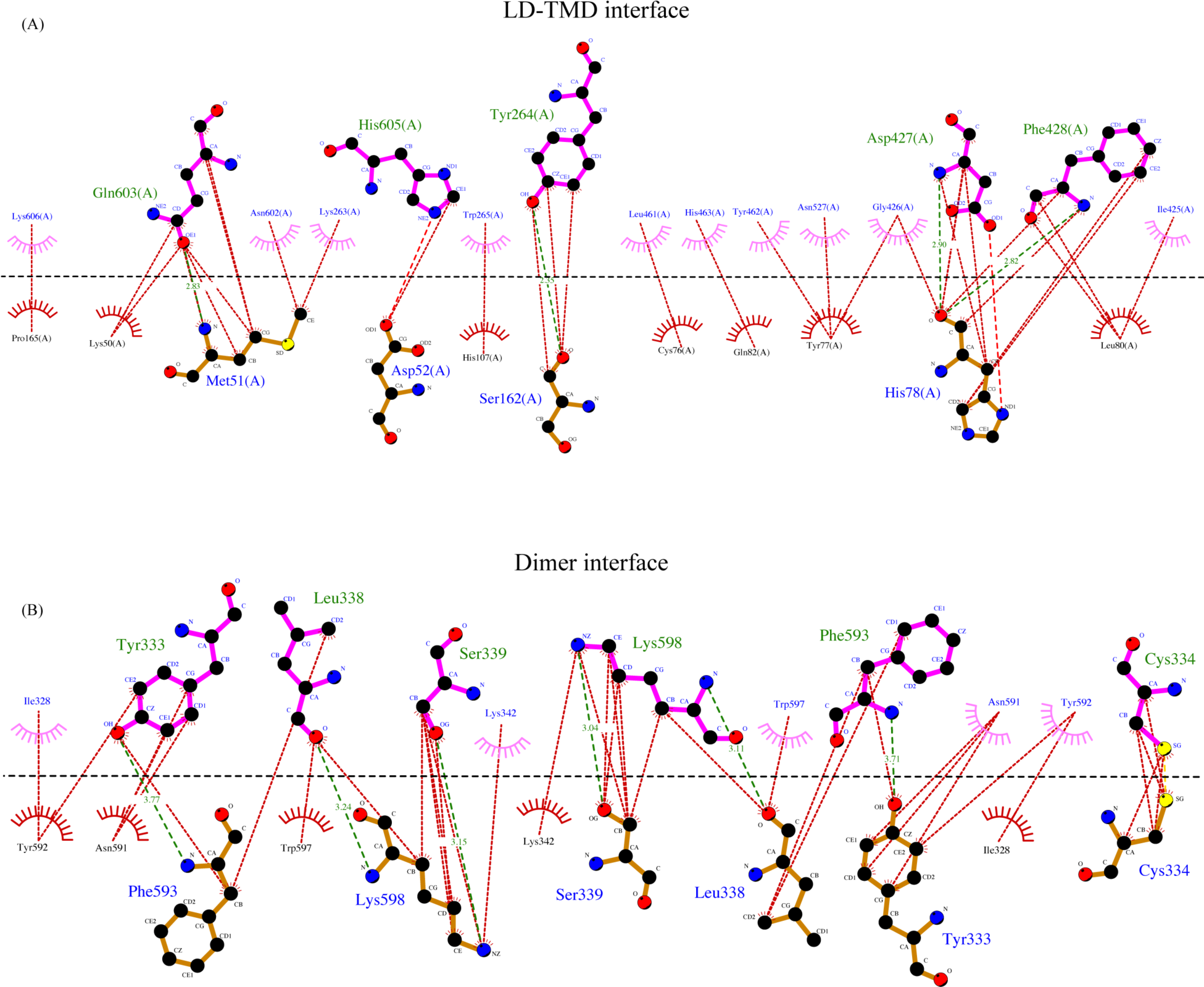
Interactions at LD-TMD and dimer interface. A 2D depiction of network of interactions (<4.5 Å) between residues at the LD-TMD interface (top, **A**) and at the dimer interface (bottom, **B**) generated in LigPlot+. Hydrogen bonds are shown as dashed lines with bond distance. Nonbonded and hydrophobic interactions are shown as dotted lines. Residues involved in nonbonded interactions are displayed as eye lashes. Chain A and chain B residues are shown in blue and orange. Dashed black line indicates the interface. The sulfurs of involved in disulfide bond at the dimer interface are highlighted in yellow.

**Figure S7:**
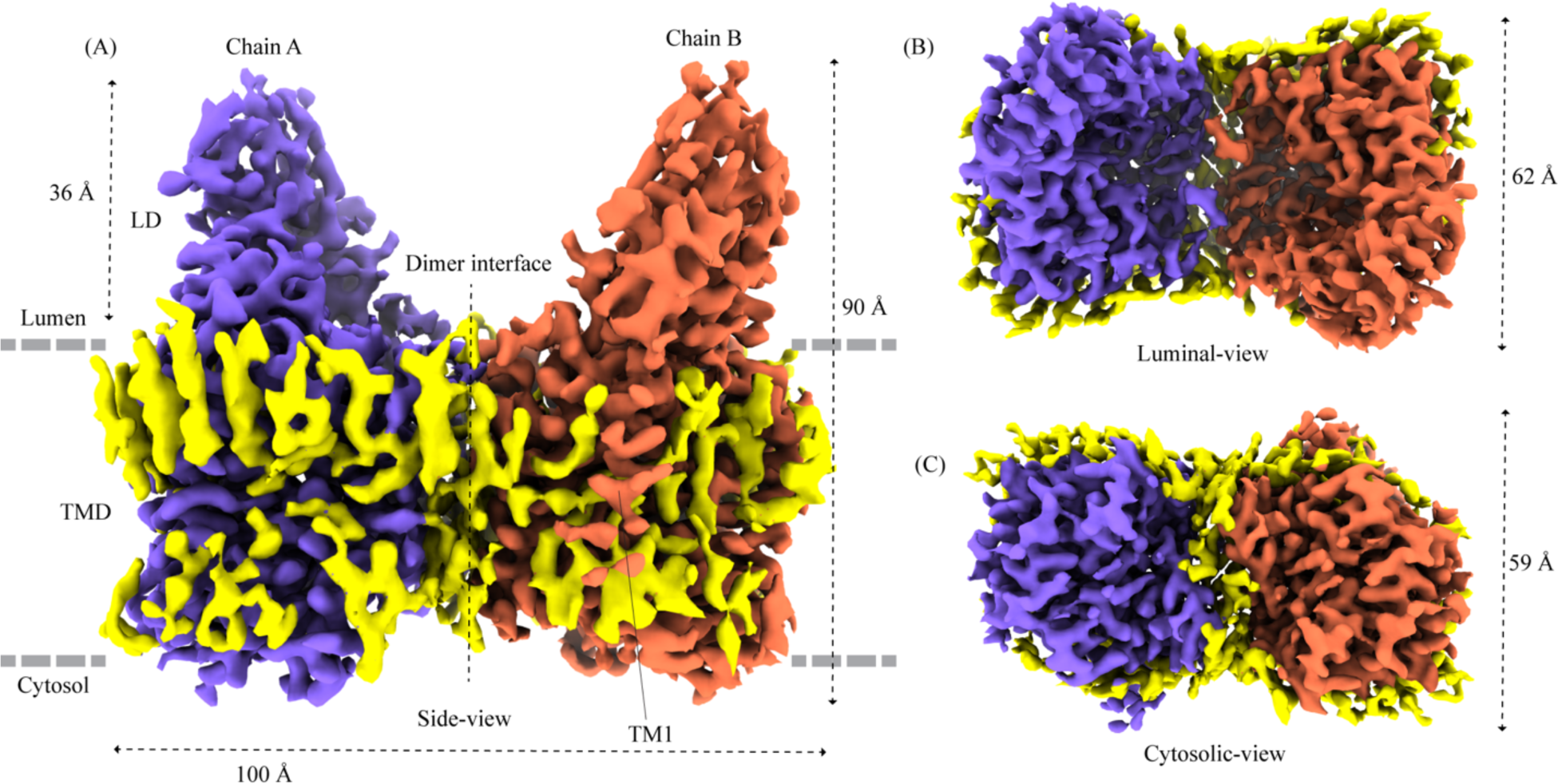
Lipids and detergent in the structure. Ordered density observed in our final cryo-EM map that did not account for protein and ligand has been displayed as yellow density (display level 0.22 of the C2 refine map in ChimeraX) in side-view **(A)**, luminal-view **(B)**, and cytosolic-view **(C)**. We believe these are ordered lipids and detergent molecules that interact with hydrophobic patches of the protein. Towards the cytosolic side (C) we find lipid/detergent density between the two protomers, forming a partition between two ACOSs. Chain A and chain B are shown in purple and orange. Dashed line indicates dimer interface.

**Figure S8:**
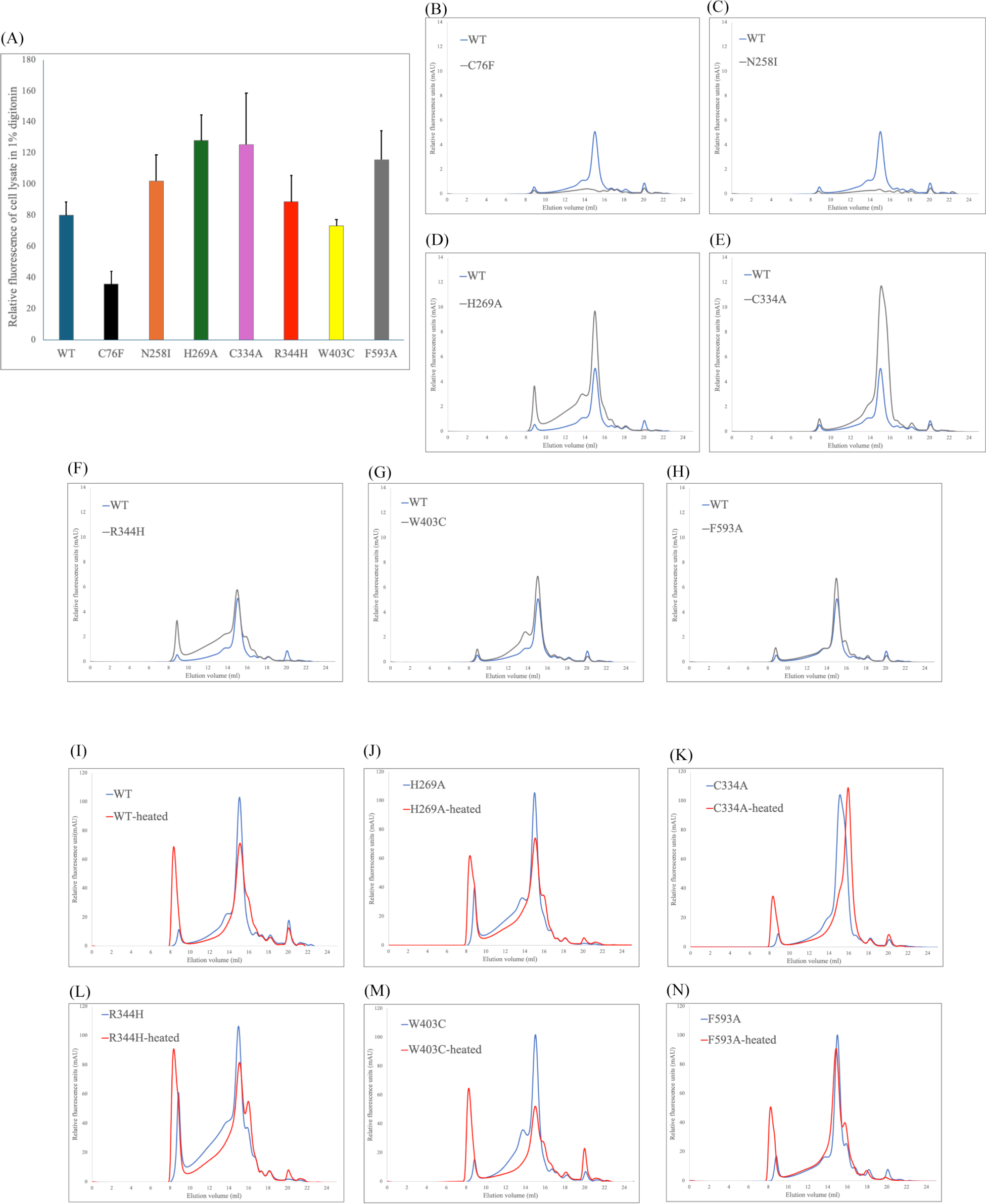
Expression and stability of HGSNAT mutants. **(A)** A comparison of relative protein expression indicated by total GFP fluorescence in 100,000 HEK293S GnTI-cells expressing HGSNAT and its mutants. **(B-H)** A comparison of FSEC chromatograms of the HGSNAT mutants (gray chromatograms) with WT HGSNAT (blue chromatogram). C76F and N258I mutants show no peak at HGSNAT dimer position, and the remaining mutants’ peak position is same as dimeric WT HGSNAT. **(I-N)** Relative stability of HGSNAT mutants analyzed by FSEC. To estimate relative stability of mutants, the solubilized mutant cell lysates were heated at 65°C for 15 min (red chromatograms) and the loss of HGSNAT peak in the resultant chromatograms were compared with non-heated samples (blue chromatograms). C334A, the mutant which breaks the disulfide at the dimer interface, results in a monomeric HGSNAT peak upon heating, while all other mutants retain their dimeric status.

**Figure S9:**
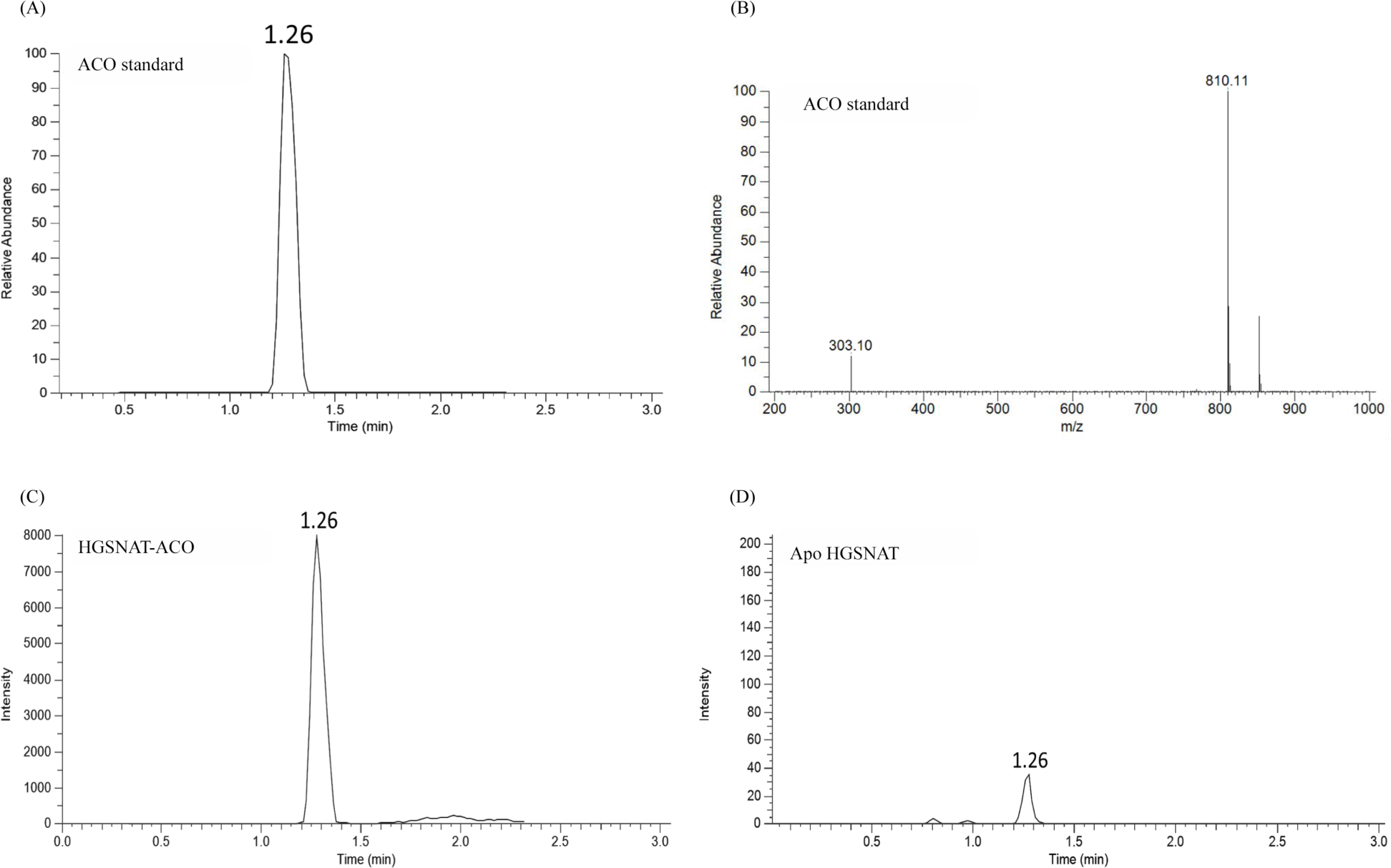
LC-MS analysis of purified HGSNAT. **(A)** LC profile and **(B)** MS/MS spectrum of acetyl-CoA (ACO) standard, showing the retention time (1.26 min), and precursor (810.1 m/z) and product (303.1 m/z) peaks in single reaction monitoring mode, respectively. **(C)** and **(D)** show relative LC peak intensities of endogenously bound ACO identified in purified HGSNAT before and after dialysis of the membranes respectively.

### Supplementary tables and legends

**Table S1:** Homologs of TMD and LD of HGSNAT found in a Dali search. Dali (**D**istance M**a**trix A**li**gnment) web server was used to search the existing database of known structures to find homologs of HGSNAT (Holm *et al,* 2023). The poor % identity (<20%) and low % sequence alignment suggests that there are no available structures of homologs of HGSNAT. Low mean RMSD of hits obtained using LD of HGSNAT as input suggests that LD is like some of the existing β-sandwiches, but TMD of HGSNAT is a novel fold.

| PDB ID | RMSD (Å) | Number of equivalent residues | Total number of residues | % Identity | Protein |
| --- | --- | --- | --- | --- | --- |
| *Homologs of TMD* |  |  |  |  |  |
| 5LNK | 5.8 | 84 | 175 | 17 | Mitochondrial complex 1, 51 kDa subunit |
| 5CE3 | 4.3 | 93 | 598 | 16 | Actin |
| 6B2Z | 9.1 | 87 | 249 | 15 | Mitochondrial ATP synthase subunit C |
| 6HV6 | 4 | 66 | 293 | 15 | PatoxP toxin |
| 6KKK | 5.9 | 133 | 380 | 14 | Sugar efflux transporter SotB |
| *Homologs of LD* |  |  |  |  |  |
| 6CZT | 2 | 81 | 82 | 15 | AlgF, alginate biosynthesis protein |
| 6IXH | 3.1 | 87 | 123 | 14 | Type VI secretion system core complex TSSJ |
| 6G7G | 3.1 | 88 | 115 | 13 | S-protein homolog (SPH) 15 |
| 5A9I | 2.3 | 95 | 194 | 12 | ECD of PepT2 |
| 5AMT | 3.7 | 84 | 105 | 12 | Intracellular growth locus E (IglE) |

**Table S2:** List of HGSNAT mutations implicated in MPS IIIC. FoldX web server was used to predict relative mutant stability (Schymkowitz *et al,* 2005). Positive total energy value indicates destabilization, with greater values meaning lower stability. Nonsense mutations indicated in the figure 4 as black were not included in FoldX calculations. Polymorphisms are italicized. All other mutants listed are missense mutations (Canals *et al,* 2011; Fan *et al,* 2006; Fedele & Hopwood, 2010; Feldhammer *et al,* 2009a; Feldhammer *et al,* 2009b; Hrebicek *et al,* 2006; Huizing & Gahl, 2020).

| **Mutation** | **Total energy (kcal/mol)** | **Region of the protein** |
| --- | --- | --- |
| G423W | 41.0 | LD-TMD interface |
| G424V | 18.8 |  |
| G424S | 11.8 |  |
| G262R | 7.5 |  |
| C76F | 5.0 |  |
| N273K | -0.6 |  |
| G133A | -1.4 |  |
| P283L | 3.0 | Catalytic core |
| R344H | 2.0 |  |
| R344C | 2.0 |  |
| E471K | 1.4 |  |
| N258I | -0.1 |  |
| G486E | 17.9 | Scaffold domain |
| M482K | 3.2 |  |
| S518F | 2.5 |  |
| W403C | 2.2 |  |
| A489E | 1.1 |  |
| ***K523Q*** | 0.9 |  |
| S539C | 0.7 |  |
| S541L | -0.2 |  |
| ***V481L*** | -0.3 |  |
| L445P | 5.9 | Other regions |
| L113P | 4.9 |  |
| L137P | 3.4 |  |
| A54V | 2.9 |  |
| ***A615T*** | 2.3 |  |
| Y627C | 1.4 |  |
| D562V | 1.2 |  |
| ***P237Q*** | 0.1 |  |
| P571L | -0.2 |  |
| G173D | -0.4 |  |
| G173E | -0.9 |  |

## Notes

### Competing Interest Statement

The authors have declared no competing interest.

### Summary of Updates

The article has been edited based on comments from the referees. Additional analysis (Figure S 8,9) has also been included.

## References

Ashkenazy H, Abadi S, Martz E, Chay O, Mayrose I, Pupko T, Ben-Tal N (2016) ConSurf 2016: an improved methodology to estimate and visualize evolutionary conservation in macromolecules. Nucleic Acids Res 44: W344–350

Ausseil J, Landry K, Seyrantepe V, Trudel S, Mazur A, Lapointe F, Pshezhetsky AV (2006) An acetylated 120-kDa lysosomal transmembrane protein is absent from mucopolysaccharidosis IIIC fibroblasts: a candidate molecule for MPS IIIC. Mol Genet Metab 87: 22–31

Bame KJ, Rome LH (1985) Acetyl coenzyme A: alpha-glucosaminide N-acetyltransferase. Evidence for a transmembrane acetylation mechanism. J Biol Chem 260: 11293–11299

Bame KJ, Rome LH (1986a) Acetyl-coenzyme A:alpha-glucosaminide N-acetyltransferase. Evidence for an active site histidine residue. J Biol Chem 261: 10127–10132

Bame KJ, Rome LH (1986b) Genetic evidence for transmembrane acetylation by lysosomes. Science 233: 1087–1089

Batzios SP, Zafeiriou DI, Vargiami E, Karakiulakis G, Papakonstantinou E (2012) Differential expression of matrix metalloproteinases in the serum of patients with mucopolysaccharidoses. JIMD Rep 3: 59–66

Beale JH, Parker JL, Samsudin F, Barrett AL, Senan A, Bird LE, Scott D, Owens RJ, Sansom MSP, Tucker SJ et al (2015) Crystal Structures of the Extracellular Domain from PepT1 and PepT2 Provide Novel Insights into Mammalian Peptide Transport. Structure 23: 1889–1899

Bonifacino JS, Traub LM (2003) Signals for sorting of transmembrane proteins to endosomes and lysosomes. Annu Rev Biochem 72: 395–447

Bork P, Holm L, Sander C (1994) The immunoglobulin fold. Structural classification, sequence patterns and common core. J Mol Biol 242: 309–320

Canals I, Elalaoui SC, Pineda M, Delgadillo V, Szlago M, Jaouad IC, Sefiani A, Chabas A, Coll MJ, Grinberg D et al (2011) Molecular analysis of Sanfilippo syndrome type C in Spain: seven novel HGSNAT mutations and characterization of the mutant alleles. Clin Genet 80: 367–374

Chidyausiku TM, Mendes SR, Klima JC, Nadal M, Eckhard U, Roel-Touris J, Houliston S, Guevara T, Haddox HK, Moyer A et al (2022) De novo design of immunoglobulin-like domains. Nat Commun 13: 5661

Coutinho MF, Lacerda L, Alves S (2012) Glycosaminoglycan storage disorders: a review. Biochem Res Int 2012: 471325

Durand S, Feldhammer M, Bonneil E, Thibault P, Pshezhetsky AV (2010) Analysis of the biogenesis of heparan sulfate acetyl-CoA:alpha-glucosaminide N-acetyltransferase provides insights into the mechanism underlying its complete deficiency in mucopolysaccharidosis IIIC. J Biol Chem 285: 31233-31242

Emsley P, Lohkamp B, Scott WG, Cowtan K (2010) Features and development of Coot. Acta Crystallogr D Biol Crystallogr 66: 486–501

Fan X, Tkachyova I, Sinha A, Rigat B, Mahuran D (2011) Characterization of the biosynthesis, processing and kinetic mechanism of action of the enzyme deficient in mucopolysaccharidosis IIIC. PLoS One 6: e24951

Fan X, Zhang H, Zhang S, Bagshaw RD, Tropak MB, Callahan JW, Mahuran DJ (2006) Identification of the gene encoding the enzyme deficient in mucopolysaccharidosis IIIC (Sanfilippo disease type C). Am J Hum Genet 79: 738–744

Fedele AO, Hopwood JJ (2010) Functional analysis of the HGSNAT gene in patients with mucopolysaccharidosis IIIC (Sanfilippo C Syndrome). Hum Mutat 31: E1574–1586

Feldhammer M, Durand S, Mrazova L, Boucher RM, Laframboise R, Steinfeld R, Wraith JE, Michelakakis H, van Diggelen OP, Hrebicek M et al (2009a) Sanfilippo syndrome type C: mutation spectrum in the heparan sulfate acetyl-CoA: alpha-glucosaminide N-acetyltransferase (HGSNAT) gene. Hum Mutat 30: 918–925

Feldhammer M, Durand S, Pshezhetsky AV (2009b) Protein misfolding as an underlying molecular defect in mucopolysaccharidosis III type C. PLoS One 4: e7434

Felisberto-Rodrigues C, Durand E, Aschtgen MS, Blangy S, Ortiz-Lombardia M, Douzi B, Cambillau C, Cascales E (2011) Towards a structural comprehension of bacterial type VI secretion systems: characterization of the TssJ-TssM complex of an Escherichia coli pathovar. PLoS Pathog 7: e1002386

Goehring A, Lee CH, Wang KH, Michel JC, Claxton DP, Baconguis I, Althoff T, Fischer S, Garcia KC, Gouaux E (2014) Screening and large-scale expression of membrane proteins in mammalian cells for structural studies. Nat Protoc 9: 2574–2585

Hirano Y, Gao YG, Stephenson DJ, Vu NT, Malinina L, Simanshu DK, Chalfant CE, Patel DJ, Brown RE (2019) Structural basis of phosphatidylcholine recognition by the C2-domain of cytosolic phospholipase A(2)alpha. Elife 8

Holm L, Laiho A, Toronen P, Salgado M (2023) DALI shines a light on remote homologs: One hundred discoveries. Protein Sci 32: e4519

Hornberg A, Eneqvist T, Olofsson A, Lundgren E, Sauer-Eriksson AE (2000) A comparative analysis of 23 structures of the amyloidogenic protein transthyretin. J Mol Biol 302: 649–669

Hrebicek M, Mrazova L, Seyrantepe V, Durand S, Roslin NM, Noskova L, Hartmannova H, Ivanek R, Cizkova A, Poupetova H et al (2006) Mutations in TMEM76* cause mucopolysaccharidosis IIIC (Sanfilippo C syndrome). Am J Hum Genet 79: 807–819

Huizing M, Gahl WA (2020) Inherited disorders of lysosomal membrane transporters. Biochim Biophys Acta Biomembr 1862: 183336

Jamali K, Kall L, Zhang R, Brown A, Kimanius D, Scheres SHW (2023) Automated model building and protein identification in cryo-EM maps. bioRxiv

Javadpour MM, Eilers M, Groesbeek M, Smith SO (1999) Helix packing in polytopic membrane proteins: role of glycine in transmembrane helix association. Biophys J 77: 1609–1618

Jimenez J, Doerr S, Martinez-Rosell G, Rose AS, De Fabritiis G (2017) DeepSite: protein-binding site predictor using 3D-convolutional neural networks. Bioinformatics 33: 3036–3042

Jobin PG, Butler GS, Overall CM (2017) New intracellular activities of matrix metalloproteinases shine in the moonlight. Biochim Biophys Acta Mol Cell Res 1864: 2043–2055

Jumper J, Evans R, Pritzel A, Green T, Figurnov M, Ronneberger O, Tunyasuvunakool K, Bates R, Zidek A, Potapenko A et al (2021) Highly accurate protein structure prediction with AlphaFold. Nature 596: 583–589

Kawate T, Gouaux E (2006) Fluorescence-detection size-exclusion chromatography for precrystallization screening of integral membrane proteins. Structure 14: 673–681

Klein U, Kresse H, von Figura K (1978) Sanfilippo syndrome type C: deficiency of acetyl-CoA:alpha-glucosaminide N-acetyltransferase in skin fibroblasts. Proc Natl Acad Sci U S A 75: 5185–5189

Koch I, Kaden F, Selbig J (1992) Analysis of protein sheet topologies by graph theoretical methods. Proteins 12: 314–323

Kretsinger RH, Kretsinger RH, Uversky VN, Permiakov EA, SpringerLink (2013) Encyclopedia of metalloproteins. Springer, New York

Kucukelbir A, Sigworth FJ, Tagare HD (2014) Quantifying the local resolution of cryo-EM density maps. Nat Methods 11: 63–65

Li F, Leier A, Liu Q, Wang Y, Xiang D, Akutsu T, Webb GI, Smith AI, Marquez-Lago T, Li J et al (2020) Procleave: Predicting Protease-specific Substrate Cleavage Sites by Combining Sequence and Structural Information. Genomics Proteomics Bioinformatics 18: 52–64

Liebschner D, Afonine PV, Baker ML, Bunkoczi G, Chen VB, Croll TI, Hintze B, Hung LW, Jain S, McCoy AJ et al (2019) Macromolecular structure determination using X-rays, neutrons and electrons: recent developments in Phenix. Acta Crystallogr D Struct Biol 75: 861–877

McBride KL, Flanigan KM (2021) Update in the Mucopolysaccharidoses. Semin Pediatr Neurol 37: 100874

Meikle PJ, Whittle AM, Hopwood JJ (1995) Human acetyl-coenzyme A:alpha-glucosaminide N-acetyltransferase. Kinetic characterization and mechanistic interpretation. Biochem J 308 (Pt 1): 327–333

Meng EC, Goddard TD, Pettersen EF, Couch GS, Pearson ZJ, Morris JH, Ferrin TE (2023) UCSF ChimeraX: Tools for Structure Building and Analysis. Protein Sci: e4792

Mirdita M, Schutze K, Moriwaki Y, Heo L, Ovchinnikov S, Steinegger M (2022) ColabFold: making protein folding accessible to all. Nat Methods 19: 679–682

Muller S, Dennemarker J, Reinheckel T (2012) Specific functions of lysosomal proteases in endocytic and autophagic pathways. Biochim Biophys Acta 1824: 34–43

Nagel L, Oliveira R, Pshezhetsky AV, Morales CR (2019) HGSNAT enzyme deficiency results in accumulation of heparan sulfate in podocytes and basement membranes. Histol Histopathol 34: 1377–1385

Newman KE, Tindall SN, Mader SL, Khalid S, Thomas GH, Van Der Woude MW (2023) A novel fold for acyltransferase-3 (AT3) proteins provides a framework for transmembrane acyl-group transfer. Elife 12

Orwick-Rydmark M, Arnold T, Linke D (2016) The Use of Detergents to Purify Membrane Proteins. Curr Protoc Protein Sci 84: 4 8 1-4 8 35

Pan X, Taherzadeh M, Bose P, Heon-Roberts R, Nguyen ALA, Xu T, Para C, Yamanaka Y, Priestman DA, Platt FM et al (2022) Glucosamine amends CNS pathology in mucopolysaccharidosis IIIC mouse expressing misfolded HGSNAT. J Exp Med 219

Parker JL, Deme JC, Wu Z, Kuteyi G, Huo J, Owens RJ, Biggin PC, Lea SM, Newstead S (2021) Cryo-EM structure of PepT2 reveals structural basis for proton-coupled peptide and prodrug transport in mammals. Sci Adv 7

Platt FM, d’Azzo A, Davidson BL, Neufeld EF, Tifft CJ (2018) Lysosomal storage diseases. Nat Rev Dis Primers 4: 27

Pravda L, Sehnal D, Tousek D, Navratilova V, Bazgier V, Berka K, Svobodova Varekova R, Koca J, Otyepka M (2018) MOLEonline: a web-based tool for analyzing channels, tunnels and pores (2018 update). Nucleic Acids Res 46: W368–W373

Pshezhetsky AV, Martins C, Ashmarina M (2018) Sanfilippo type C disease: pathogenic mechanism and potential therapeutic applications. Expert Opin Orphan D 6: 635–646

Punjani A, Rubinstein JL, Fleet DJ, Brubaker MA (2017) cryoSPARC: algorithms for rapid unsupervised cryo-EM structure determination. Nat Methods 14: 290–296

Punjani A, Zhang H, Fleet DJ (2020) Non-uniform refinement: adaptive regularization improves single-particle cryo-EM reconstruction. Nat Methods 17: 1214–1221

Rajasekar KV, Ji S, Coulthard RJ, Ride JP, Reynolds GL, Winn PJ, Wheeler MJ, Hyde EI, Smith LJ (2019) Structure of SPH (self-incompatibility protein homologue) proteins: a widespread family of small, highly stable, secreted proteins. Biochem J 476: 809–826

Rome LH, Crain LR (1981) Degradation of mucopolysaccharide in intact isolated lysosomes. J Biol Chem 256: 10763–10768

Rome LH, Hill DF, Bame KJ, Crain LR (1983) Utilization of exogenously added acetyl coenzyme A by intact isolated lysosomes. J Biol Chem 258: 3006–3011

Rudnik S, Damme M (2021) The lysosomal membrane-export of metabolites and beyond. FEBS J 288: 4168–4182

Ruivo R, Anne C, Sagne C, Gasnier B (2009) Molecular and cellular basis of lysosomal transmembrane protein dysfunction. Biochim Biophys Acta 1793: 636–649

Saier MH, Reddy VS, Moreno-Hagelsieb G, Hendargo KJ, Zhang Y, Iddamsetty V, Lam KJK, Tian N, Russum S, Wang J et al (2021) The Transporter Classification Database (TCDB): 2021 update. Nucleic Acids Res 49: D461–D467

Schroder B, Saftig P (2016) Intramembrane proteolysis within lysosomes. Ageing Res Rev 32: 51–64

Schwake M, Schroder B, Saftig P (2013) Lysosomal membrane proteins and their central role in physiology. Traffic 14: 739–748

Schymkowitz J, Borg J, Stricher F, Nys R, Rousseau F, Serrano L (2005) The FoldX web server: an online force field. Nucleic Acids Res 33: W382–388

Song J, Tan H, Perry AJ, Akutsu T, Webb GI, Whisstock JC, Pike RN (2012) PROSPER: an integrated feature-based tool for predicting protease substrate cleavage sites. PLoS One 7: e50300

Steenhuis P, Froemming J, Reinheckel T, Storch S (2012) Proteolytic cleavage of the disease-related lysosomal membrane glycoprotein CLN7. Biochim Biophys Acta 1822: 1617–1628

Turk B, Turk D, Turk V (2000) Lysosomal cysteine proteases: more than scavengers. Biochim Biophys Acta 1477: 98–111

Ulmschneider MB, Sansom MS (2001) Amino acid distributions in integral membrane protein structures. Biochim Biophys Acta 1512: 1–14

Varadi M, Anyango S, Deshpande M, Nair S, Natassia C, Yordanova G, Yuan D, Stroe O, Wood G, Laydon A et al (2022) AlphaFold Protein Structure Database: massively expanding the structural coverage of protein-sequence space with high-accuracy models. Nucleic Acids Res 50: D439–D444

Xu R, Ning Y, Ren F, Gu C, Zhu Z, Pan X, Pshezhetsky AV, Ge J, Yu J (2024) Structure and mechanism of lysosome transmembrane acetylation by HGSNAT. Nat Struct Mol Biol. DOI: 10.1038/s41594-024-01315-5

